# Mathematical generation of data-driven hippocampal CA1 pyramidal neurons and interneurons copies via A-GLIF models for large-scale networks covering the experimental variability range

**DOI:** 10.1101/2023.04.03.535350

**Authors:** A. Marasco, C. Tribuzi, A. Iuorio, M. Migliore

**Affiliations:** Department of Mathematics and Applications, University of Naples Federico II, Naples, Italy; Institute of Biophysics, National Research Council, Palermo, Italy; University of Vienna, Faculty of Mathematics, Vienna, Austria

**Keywords:** Neuronal modeling, adaptive generalized leaky integrate-and-fire (A-GLIF) models, hippocampal CA1 pyramidal neurons and interneurons, neuron copies, EBRAINS

## Abstract

Efficient and accurate large-scale networks are a fundamental tool in modelling brain areas, to advance our understanding of neuronal dynamics. However, their implementation faces two key issues: computational efficiency and heterogeneity. Computational efficiency is achieved using simplified neurons, whereas there are no practical solutions available to solve the problem of reproducing in a large-scale network the experimentally observed heterogeneity of the intrinsic properties of neurons. This is important, because the use of identical nodes in a network can generate artifacts which can hinder an adequate representation of the properties of a real network.

To this aim, we introduce a mathematical procedure to generate an arbitrary large number of copies of simplified hippocampal CA1 pyramidal neurons and interneurons models, which exhibit the full range of firing dynamics observed in these cells - including adapting, non-adapting and bursting. For this purpose, we rely on a recently published *adaptive generalized leaky integrate-and-fire (A-GLIF)* modeling approach, leveraging on its ability to reproduce the rich set of electrophysiological behaviours of these types of neurons under a variety of different stimulation currents.

The generation procedure is based on a perturbation of model’s parameters related to the initial data, firing block, and internal dynamics, and suitably validated against experimental data to ensure that the firing dynamics of any given cell copy remains within the experimental range. This allows to obtain heterogeneous copies with mathematically controlled firing properties. A full set of heterogeneous neurons composing the CA1 region of a rat hippocampus (approximately 500K neurons), are provided in a database freely available in the *live paper* section of the EBRAINS platform.

By adapting the underlying A-GLIF framework, it will be possible to extend the numerical approach presented here to create, in a mathematically controlled manner, an arbitrarily large number of non-identical copies of cell populations with firing properties related to other brain areas.

## 1 Introduction

Computational models of multiple brain areas provide an invaluable tool to make significant advances to understand how a brain works, from cognitive functions and dysfunctions to digital twin implementations.

Several techniques have been implemented for this purposes. One of them is to consider the neural dynamics collectively through a mean-field approach, to obtain dimensionality reduction [2, 3, 4, 28]. Another well-established strategy relies on the construction of large-scale networks linking the dynamics of individual neurons (e.g. using a morphologically and biophysically accurate implementation [6, 17, 18], LIF models [14, 31] or the Izhikevich model [8, 16]). To obtain a solid representation of a mean-field reduction it is necessary to start from a detailed understanding not only of individual neuron dynamics but also of the network to which they belong [26]. Efficient and accurate large-scale networks hence represent an essential tool to study neural dynamics. However, in dealing with such structures, two crucial issues arise: computational efficiency and heterogeneity. Computational efficiency is required to ameliorate the current technical limitations of supercomputer systems, especially in terms of energy, computational, and memory requirements. From this point of view, neuron models (and their networks) that achieve a good compromise between accuracy and computational efficiency are the generalized leaky integrate-and-fire (GLIF) models, complemented by appropriate initial and update conditions (e.g. [9, 27, 30]). Extensions of this modeling framework include the extended GLIF (E-GLIF) and the adaptive GLIF (A-GLIF), introduced by Geminiani et al. in [5] and Marasco et al. in [15], respectively. The A-GLIF framework, in particular, allows to obtain better constraints in the model parameters’ space, and to obtain a quantitative agreement with the observed number and timing of spikes experimentally observed in 84 CA1 neurons and interneurons in response to a wide range of input currents.

Cells heterogeneity is another important issue. Large-scale neural networks incorporating cell type diversity (due to e.g. intrinsic physiology, morphology, connectivity, and genetic identity) have in fact been proven to strongly influence the emergent properties of neural networks, hence playing a fundamental role in the information processing in the nervous system (see [29] and references therein). While implementing identical neurons in a network is biologically unrealistic [11], the use of non-identical neurons may be analytically and computationally more challenging, although they leads to networks which better reproduce different types of firing behaviour, including synchronisation [1, 12, 13, 14, 16, 21, 23, 24]. However, Virtually all networks of simplified neurons still use identical neurons.

Libraries of different sizes, providing non-identical neuron copies based on available experimental data, have been obtained either perturbing the intrinsic model parameters of point-neuron models (e.g. [7, 8, 20, 29] or adding noise to perturb morphological features in such a way to create copies whose firing patterns remains within experimental ranges (e.g. [10, 22, 25] and references therein). For accurate morphological and biophysical implementations (e.g. [18, 17], this problem has been solved using cloning procedures able to generate an appropriate number of cells with individual prperties consistent with the experimental variability.

These ex-post approaches do not allow to anticipate the firing behaviour of a clone, and this may significantly slow down the process of creating neurons with the proper electrophysiological properties representing, for example, specific sub-populations or distributions.

In this work, we introduce a novel methodology to implement copies representing the different firing dynamics of CA1 neurons and interneurons, by perturbing specific parameters in the A-GLIF model introduced in [15]. We show how it is possible to control the dynamical behaviour of the copies and, at the same time, to remain bounded within the experimental range.

The database of approximately 500K neurons, provided in the *live paper section of EBRAINS (https://ebrains.eu/service/live-papers/)*, represents the entire neurons population of a rat hippocampal CA1 area. The code can be easily extended to generate an arbitrary larger population, to consider other brain regions.

The paper is structured as follows: in Section 2 we introduce the reference experimental data used to obtain the database, together with the A-GLIF modelling framework used to investigate neurons’ dynamics. In Section 3 we discuss different cloning procedures and the algorithm to generate copies with controlled firing properties. We conclude our work with a discussion of the results and an outline of future research perspectives in Section 4.

## 2 Materials and Methods

### 2.1 Experimental data

As a reference to implement our procedure, we considered a set of over 500 somatic voltage traces recorded from 84 cells: 58 pyramidal and 26 interneurons, obtained from *in vitro* rat hippocampal CA1 slices [19], in response to somatic constant current injections, from 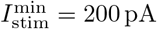 to 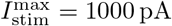. The physiological variability for pyramidal neurons and interneuron is shown in Fig. 1. The 314 traces from pyramidal neurons (Fig. 1A) were all classified as *continuous accommodating cells* (cAC); for interneurons (Fig. 1B), 54 traces were classified as cAC, 72 traces as *bursting cells* (bAC), and 62 traces as *continuous non-accommodating cells* (cNAC). To better illustrate the different firing behaviour within each class, in Fig. 2 we show typical examples of number of spikes as a function of the input current. Note the rather different results, in response to the same input, observed for different cells in both pyramidal cells (Fig. 2, left panel) and interneurons (Fig. 2, right panel).

**Figure 1:**
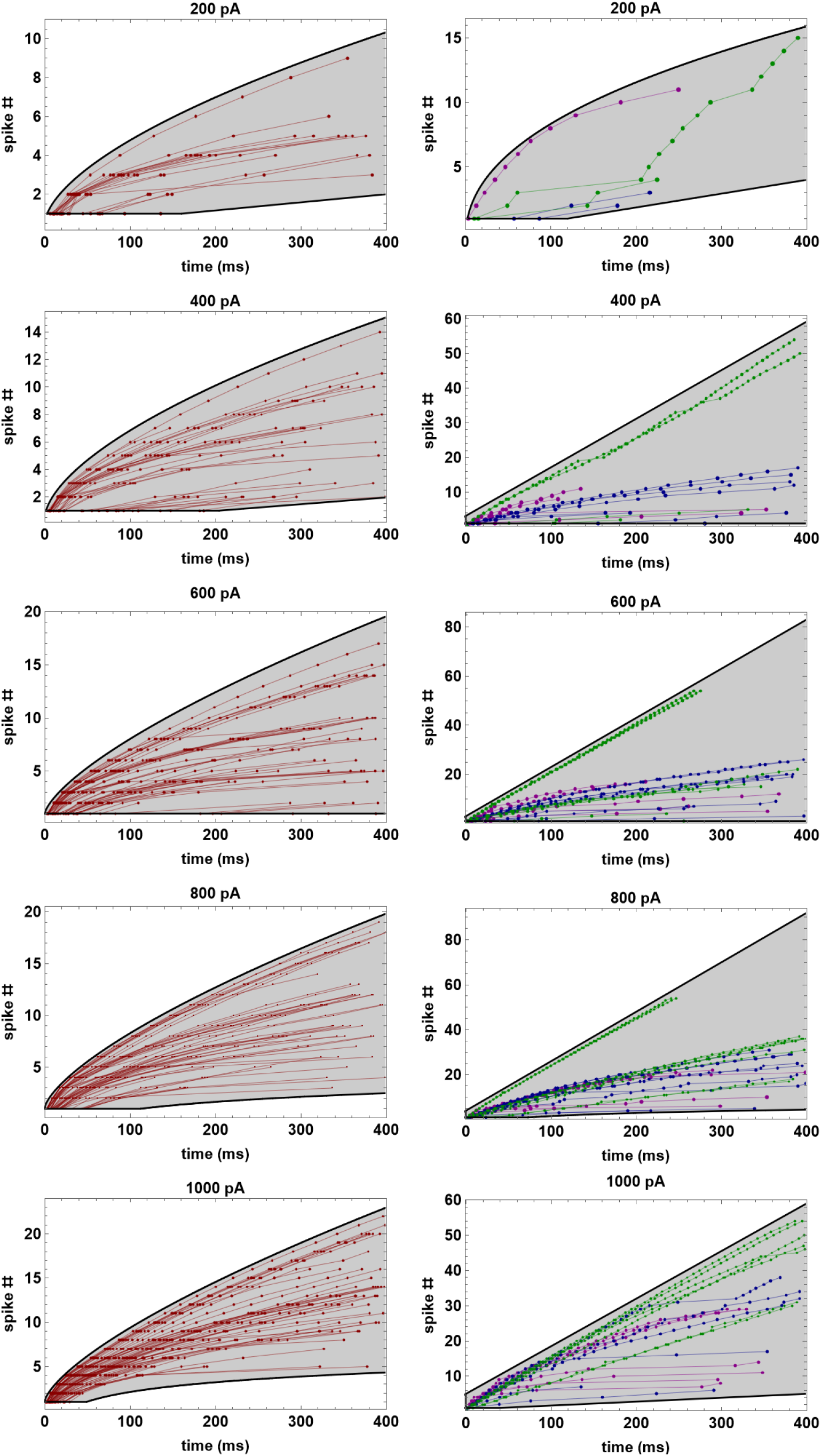
Reference experimental data and their range of variability. Spike number as a function of spike times for pyramidal neurons (left column, red markers and lines) and CA1 interneurons (right column) classified as cAC (blue markers and lines), bAC (magenta markers and lines), and cNAC (green markers and lines). Gray areas represent the variability regions bounded by the black curves (detailed expression available in the Supplementary information).

**Figure 2:**
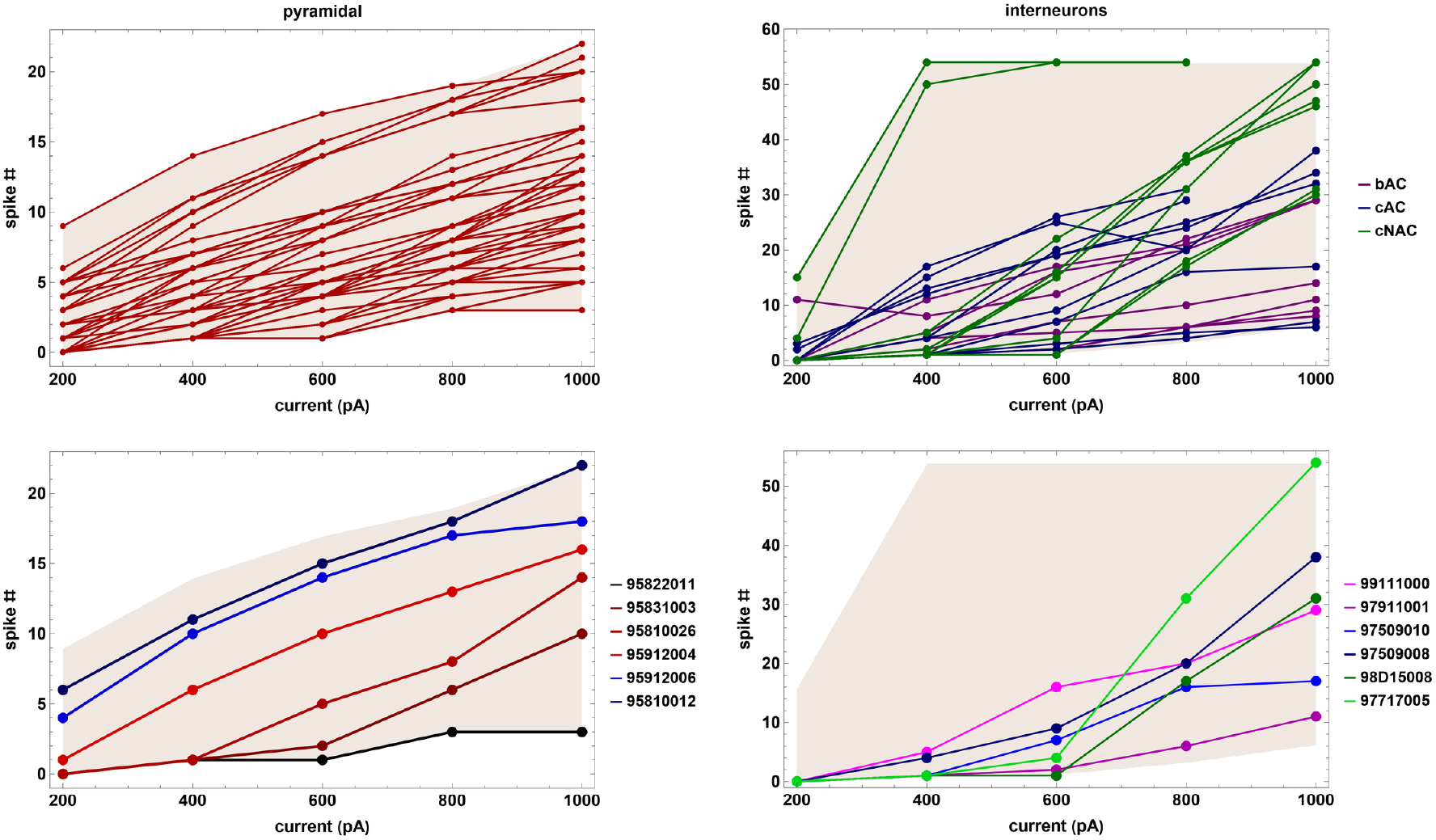
Number of spikes as a function of the stimulation current. Pyramidal neurons (left column) and interneurons (right column) of type cAC (blue markers and lines), bAC (magenta markers and lines), and cNAC (green markers and lines). The lower panels correspond to typical data for the spike number as a function of the stimulation currents for some pyramidal neurons (left) and interneurons (right).

### 2.2 The A–GLIF model

#### 2.2.1 Model equations, initial and update conditions

In [15] we introduced the A-GLIF model which aims to describe the evolution, in a subthreshold regime, of the membrane potential *V* coupled with the adaptation (*I*_adap_) and depolarization (*I*_dep_) currents as follows:

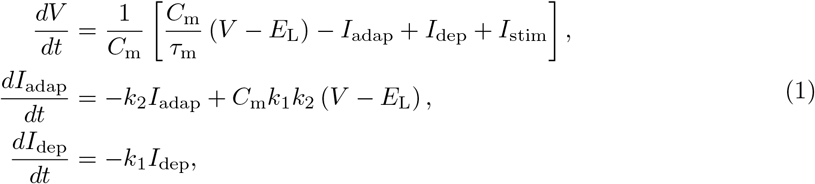

where all the parameters are positive, except for the resting potential *E_L_*, and the injected current *I*_stim_. Their explanation is given in Table 1.

**Table 1:** List of parameters appearing in Eq. (1) including their description and measurement unit.

| Parameter | Description | UM |
| --- | --- | --- |
| $E_L$ | resting potential | mV |
| $V_r$ | reset potential | mV |
| $V_{th}$ | threshold potential | mV |
| $\Delta t_{ref}$ | refractory interval | ms |
| $I_{stim}$ | external stimulation current | pA |
| $C_m$ | membrane capacitance | pF |
| $\tau_m$ | membrane time constant | ms |
| $I_{th}$ | threshold stimulation current | pA |
| $k_2$ | $I_{adap}$ decay rate | $ms^{-1}$ |
| $k_1$ | $I_{dep}$ decay rate | $ms^{-1}$ |

We assume that the neuron is at rest – i.e., *I*_stim_ = 0 and *V* = *E_L_* – for *t* < *t*_start_, where *t*_start_ represents the first time instant at which the stimulation current is different from zero. Moreover, we denote with *I*_th_ the *threshold current* above which the neuron starts to fire, i.e. we assume that a spike event occurs when, for *I*_stim_ > *I*_th_, the potential *V* reaches the *threshold potential V*_th_.

Starting from the resting condition, the *first spike* for any *I*_stim_ > *I*_th_ can be obtained by setting the initial conditions of the Cauchy problem associated to system (1) as follows

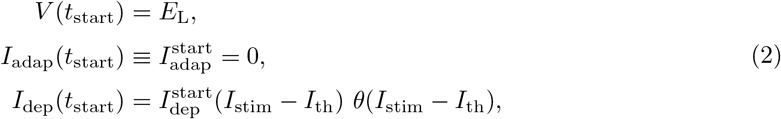

where 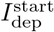 is a suitable constant and *θ*(*I*_stim_ – *I*_th_) is the step function defined as

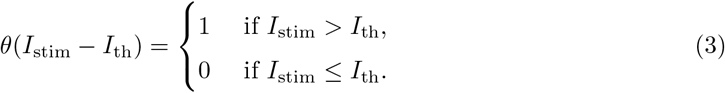

Coherently with the LIF framework, the potential *V* after a spike does not return to the resting value *E*_L_ but at the *reset potential V*_r_. Then, for any following spike, the initial conditions of each Cauchy problem associated to (1) are modified according to the following *after-spike update rules*

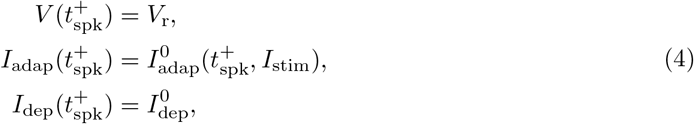

where 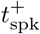 is the time instant following the spike time *t*_spk_, i.e. 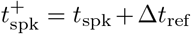, with Δ*t*_ref_ = 2 ms defined as the *refractory time*, 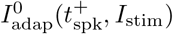 is a suitable set of initial values that depend on both the stimulation current *I*_stim_ and the corresponding spike times, and 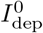 is a constant.

#### 2.2.2 Nondimensional formulation of the model

Considering the following rescaled variables

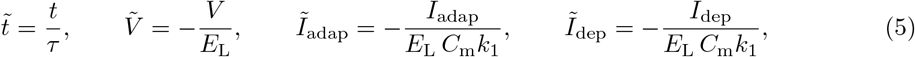

we obtain an equivalent nondimensional version of system (1) (for simplicity, from now on we will omit the tildes)

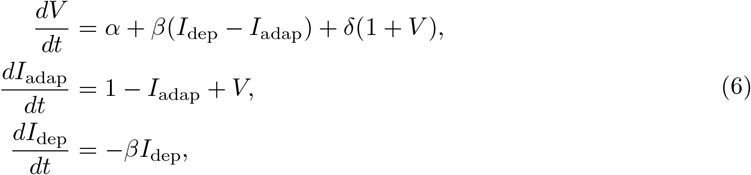

where

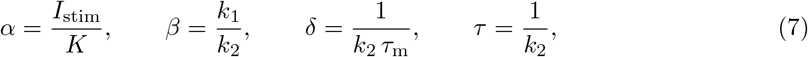

and *K* = –*C_m_ E_L_ k*_2_ is a positive scaling constant.

Similarly, the dimensionless initial conditions (2) and (4) assume the following forms, respectively,

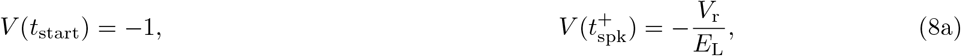

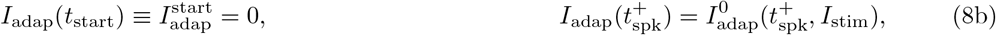

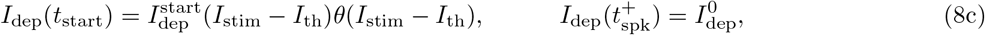

where all time variables have been rescaled by means of *τ* = 1/*k*_2_.

In [15], assuming a constant stimulation current *I*_stim_, we obtained the analytical form of the solutions of the linear system (6) as follows

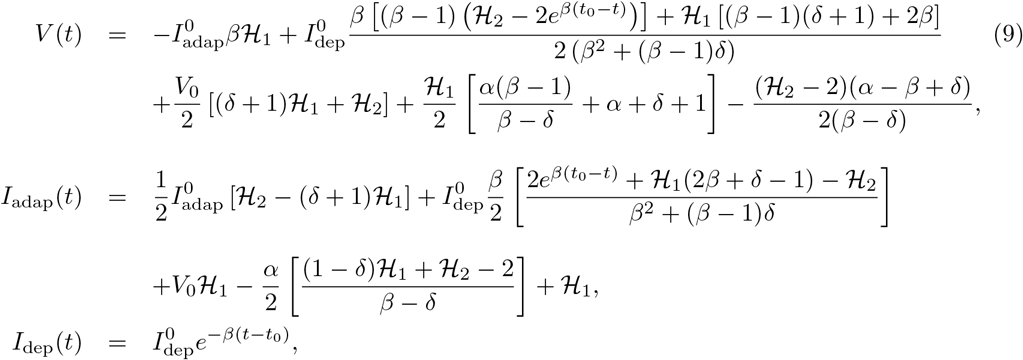

where the initial data are defined according to (8) as

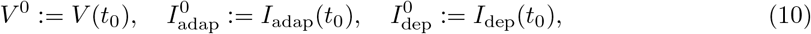

and

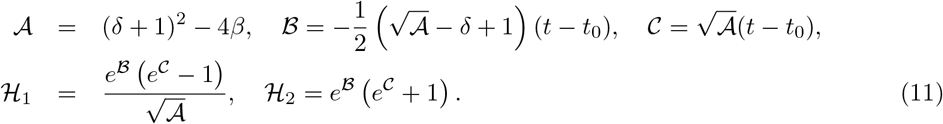

#### 2.2.3 Equilibria, parameter constraints and initial data distributions

Despite the linearity of the equations, the A-GLIF model can reproduce a wide range of physiological firing patterns like bursting, non adapting, and continuous adapting [19], provided that some parameter constraints are imposed and initial data distributions, 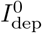 and 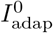, are suitably chosen. The quantitative agreement between model and experiments is shown in Fig. 3, where we compare findings for both pyramidal neurons and interneurons. Below we report the fundamental results of this analysis; an in-depth investigation can be found in [15].

**Figure 3:**
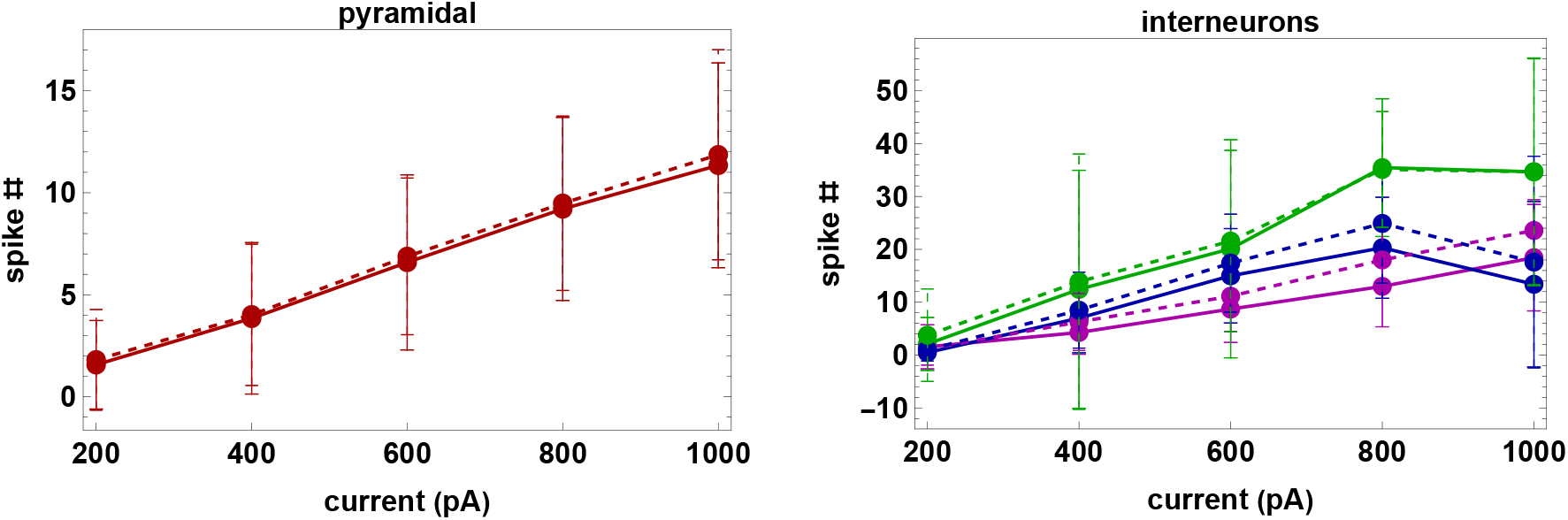
Comparison between experiments and models. Mean and standard deviation for the number of spikes as a function of the input current for pyramidal neurons (red markers and line) and interneurons cAC (blue markers and lines), bAC (magenta markers and lines), and cNAC (green markers and lines). The continuous and dashed lines represent experimental data and model results, respectively.

For *α* ≠ 0 or *β* ≠ *δ*, the dynamical system (6) admits the equilibrium

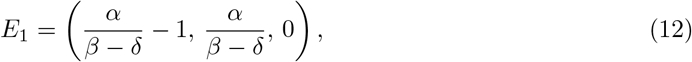

that is (globally) asymptotically stable if and only if

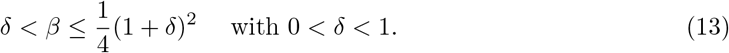

Moreover, by imposing that the cell does not fire for 0 < *I*_stim_ < *I*_th_ we obtain

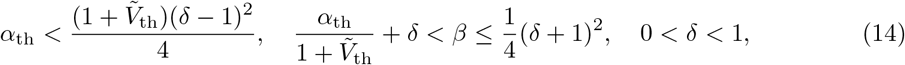

where *Ṽ*_th_ = –*V*_th_/*E*_L_ is the nondimensional form of the threshold potential *V*_th_, and *α*_th_ = *I*_th_/*K*.

Since *V*(*t*) is an increasing function for any positive stimulation current *I*_stim_, the initial data 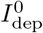 and 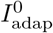 must satisfy the following condition (see Eq. (8))

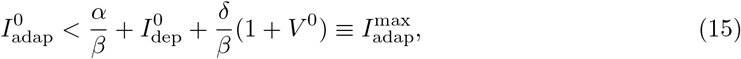

where *V*^0^ = −1 or *V*^0^ = –*V_r_*/*E_L_* for the first or after the first spike event, respectively.

In [15] we proved that the 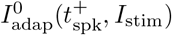 distribution for any given cell can be represented by a Monod-type function as follows^1^

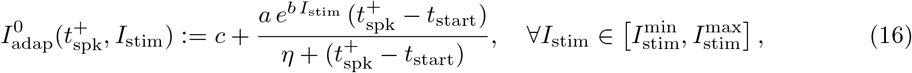

where *a*, *b*, *c*, *η* are constants, and *t*_start_ is the last instant in which *I*_stim_ = 0 or *I*_stim_ ≤ *I*_th_.

For sake of simplicity, we assume that the function (16) is defined for all 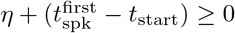, where 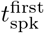 is the time of the first spike event for the current *I*_stim_. Since 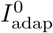 must be non-negative for all 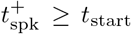, it must necessarily be *c* ≥ 0, also when *a* = 0. Moreover, Eq. (16) is an increasing monotone function of *t* when *a η* > 0 and decreasing monotone for *a η* < 0.

The plateau value of 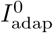, defined as

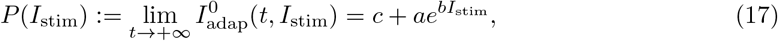

is an increasing monotone function of *I*_stim_ when *a b* > 0 and decreasing monotone when *a b* < 0.

Imposing that the function (16) is positive for any stimulation current and at any time, we obtain the following additional constraints on parameters *a*, *b*, *c*, *η*:

i. if *a*, *b* < 0 then *c* ≥ –*a* and *η* < 0;
ii. if *a* > 0, ∀*b* then *η* > 0.

The A-GLIF model generates a uniform *ISI* sequence when *a* = 0 and a nonuniform one when the parameters *a*, *b*, *c*, *η* are suitably chosen. By simultaneously fitting, for each neuron, the set of 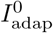 values for all experimental currents using Eq. (16) (solid curves in Fig. 4), we obtained a function which allows us to predict the spike times of a given neuron for any constant current injection in the interval 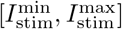.

Several experimental traces show an unexpected block of firing, long before the end of the stimulation at constant current. To reproduce these *firing blocks* for any neuron and any stimulation current I_stim_, in [15] we implemented a *Monod block procedure* that allowed us to determine the time interval in which the Monod function (16) should be defined. First, we identified the range of currents 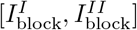 such that for 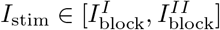 a firing block occurs, i.e.

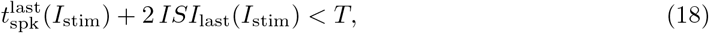

where 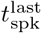 and *ISI*_last_ are the time and the *ISI* of the last spike event for the current *I*_stim_, respectively, and [*t*_start_, *T*] is the stimulation interval. Subsequently, we determined the function

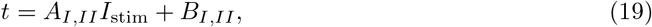

where 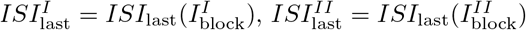, and

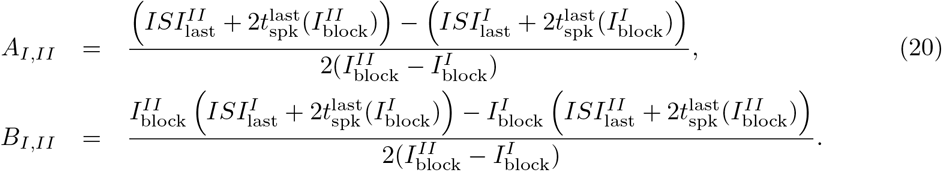

**Figure 4:**
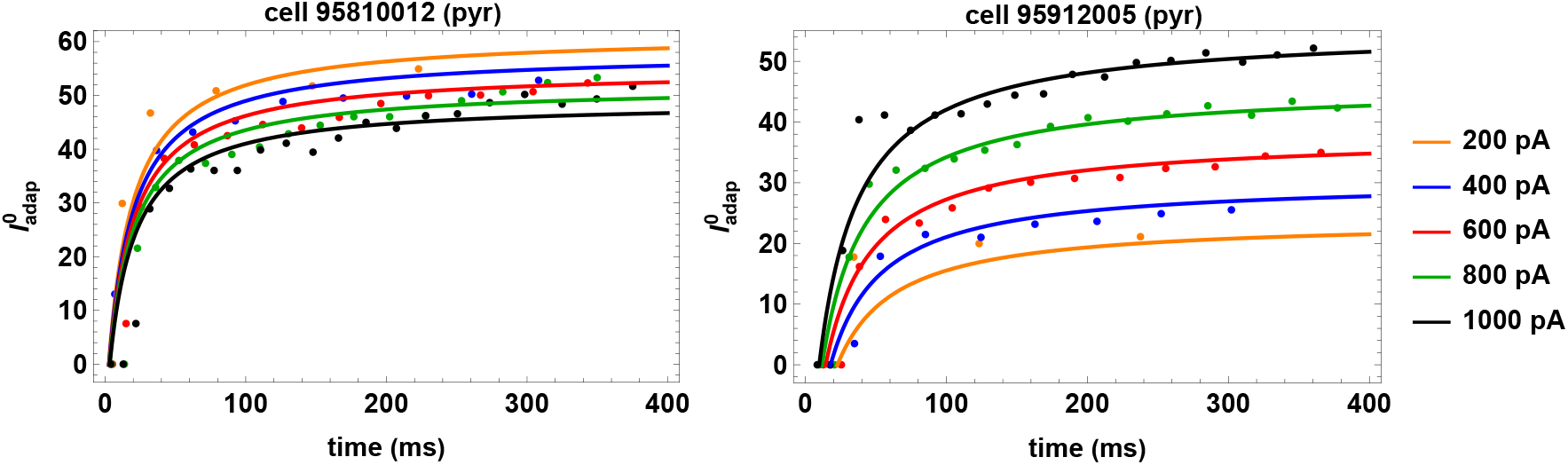
Monod-type functions. Monod functions (continuous lines) interpolating the initial data distributions 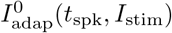 (dots) for the pyramidal cells 95810012 (left panel) and 95912005 (right panel).

Finally, denoting the stimulation current starting from which the firing block does not occur by I_fire_, the function (19) provides the time interval in which the Monod function (16) is defined as follows

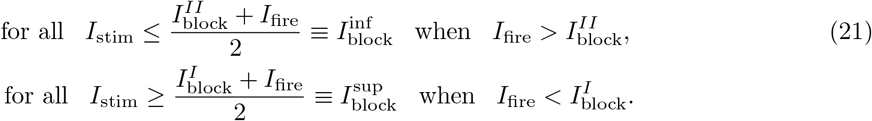

This procedure can be easily generalized when there is only one *I*_block_ value that satisfies condition (18) by setting 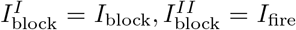 when *I*_fire_ > *I*_block_; and 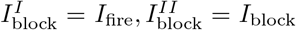 if *I*_fire_ < *I*_block_.

#### 2.2.4 Monotonicity property of *V* with respect to *α*

In this section, we show that the function *V*(*t*) provided in Eq. (9) is monotonically increasing with respect to *α*. In particular, we show that the function

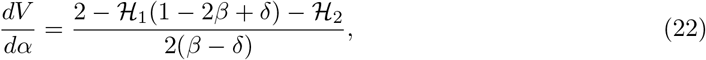

(where the functions 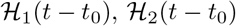 have been defined in (11)) is positive for any *t* > *t*_0_, as also numerically confirmed in Fig. 5.

**Figure 5:**
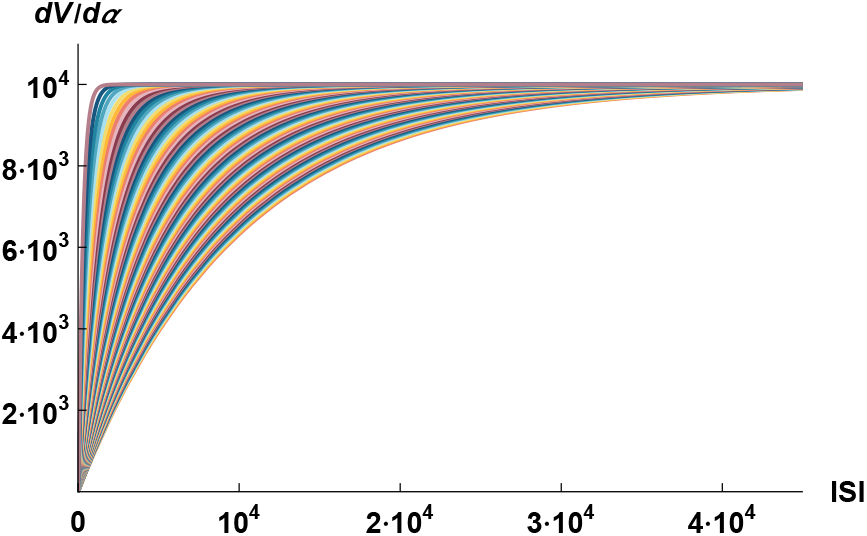
Plot of the derivative of *V* respect to *α*. Plot of the function 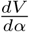 in Eq. (22) as function of *t* – *t*_0_ obtained by varying *β* and *δ* within the admissibility ranges defined by conditions (13) with a step of 0.01 for both parameters.

Equivalently, introducing *T*:= *t*–*t*_0_ we show that 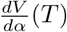 is strictly positive in *T* for any *T* > 0. This relies on the fact that:

i. 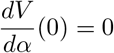
ii. 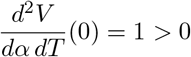
iii. 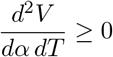 for any *β*, *δ* satisfying Eq. (13)
iv. 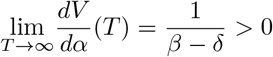 for any *β*, *δ* satisfying Eq. (13).

Conditions (i), (iv) directly follow from Eq. (22). On the other hand, conditions (ii) and (iii) are obtained by rewriting 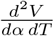 in terms of *β*, *δ*, and *T* first

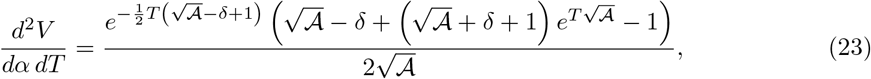

where 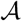 has been defined in (11).

Substituting *T* = 0 in (23) leads to condition (ii). As for (iii), we observe that the denominator of (23) is always positive under the stability conditions on *β* and *δ* given in Eq. (13). On the other hand, the numerator is positive if and only if

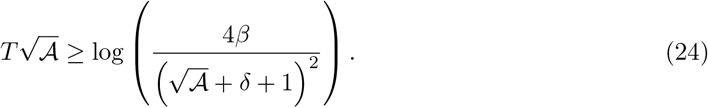

Here, the left-hand side is positive, whereas the right-hand side is negative since the argument of the logarithm is less or equal than 1 for any *β*, *δ* satisfying Eq. (13). This inequality is then automatically satisfied, leading to condition (iii).

## 3 Results

### 3.1 Mathematical procedures for models generation

The goal of this section is to describe several mathematical procedures that can be used to generate model neurons with firing properties within the experimental variability range (see Figs. 1–2).

#### 3.1.1 Temporal shift of all spike times

In this section, we discuss the procedures to delay/anticipate the first spike time and changes all other spike times. This can be achieved by modulating 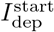 and 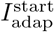 in Eqs. (8), with all the other parameters in Eqs. (7) and (16) fixed.

To this aim, it should be noted that under the constraints (13) the potential *V* is a decreasing (resp. increasing) function with respect to 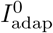 (resp. 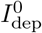) when the other initial data are fixed.

In fact, we have

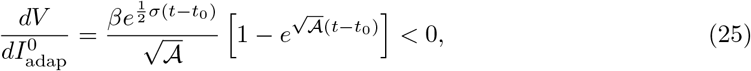

and it can be numerically proved that (see Supp. Fig. 1)

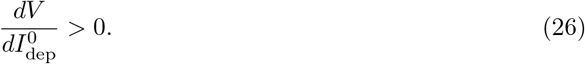

To obtain a different time of first spike, at least one of the values 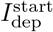 and 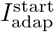 in Eqs. (8) must be varied. In view of Eqs. (16), modifying the first spike time results in a different distribution for the initial data 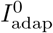, thus obtaining a different distribution of spike times (see Supp. Fig. 2 and Fig. 6).

**Figure 6:**
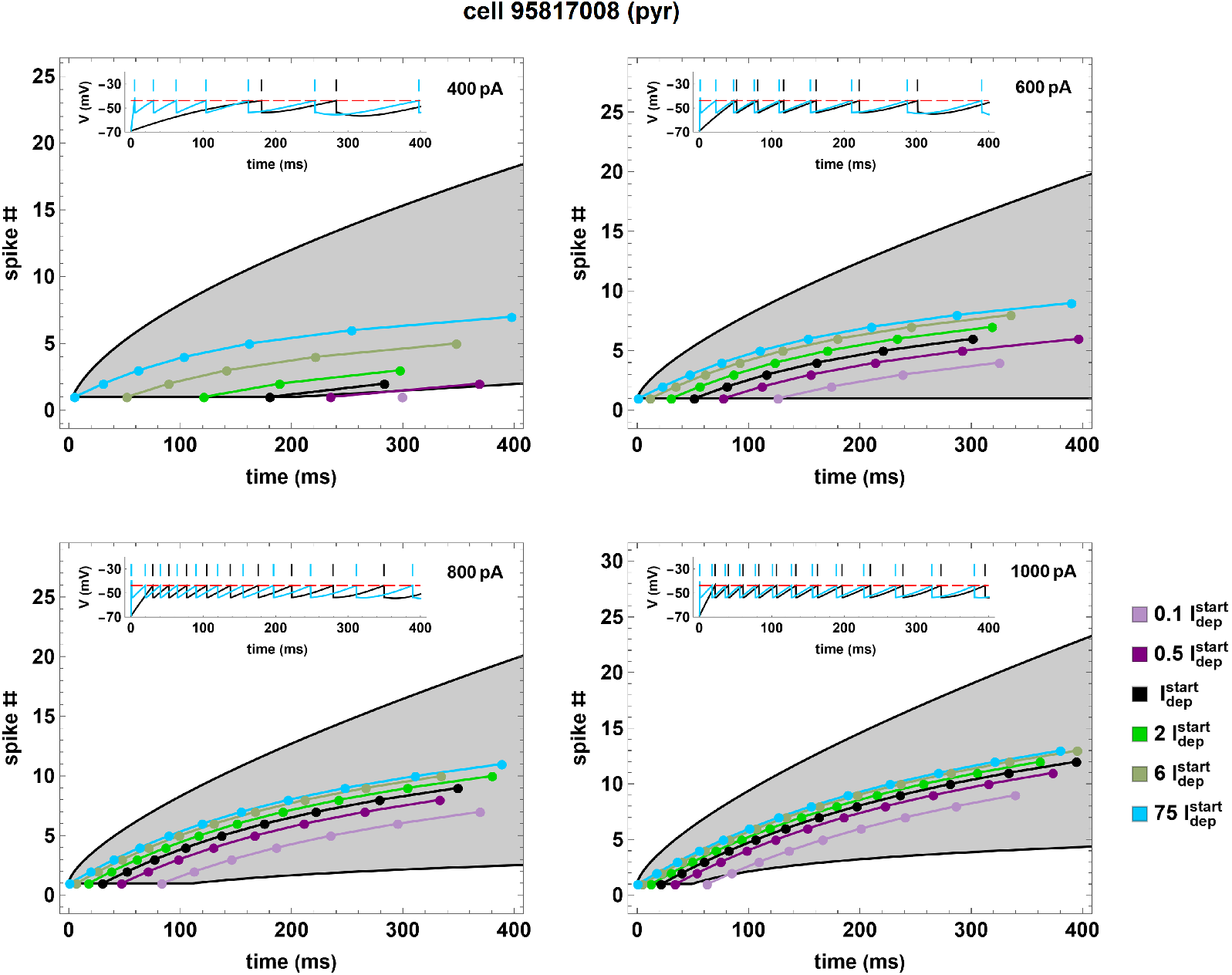
Spike number as a function time for the pyramidal cell 95817008 (black curves and dots) and for 5 neuron copies (colored curves and dots) obtained by fixing the numerical values of all parameters except 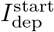. For all copies condition (15) is verified. The inset in each panel represents the original simulation compared with the copy corresponding to 75 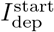.

Owing to Eqs. (16), (25), and (26), we can modulate the time of both the first and any subsequent spikes by modifying the value of 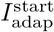 and/or 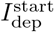. In particular, increasing the values of 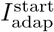 or 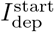 will result in increasing or decreasing first spike times, and vice versa, respectively. We note that since *I*_adap_ is a nonnegative function, in view of Eq. (8b) we can only choose positive values of 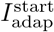. This leads to an increase in the first spike time compared to the original model (see colored curves and dots in Supp. Fig. 2). Moreover, in view of Eqs. (15) and (25), for any fixed value of 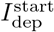, the minimum and maximum values of the first spike time can be obtained by setting the values of 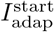, respectively, as follows

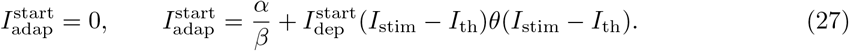

and, consequently, by setting 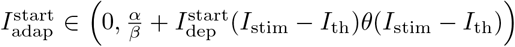 we can modify the time of all spikes including the first one (see Supp. Fig. 2).

Similarly, we can modify the time of the first spike by choosing a different value for 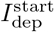 (see Fig. 6). In particular, in view of Eqs. (15) and (26), for any fixed value of 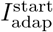, we can choose arbitrary positive values for 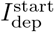 provided that condition (15) is satisfied. This leads to an early or late first spike time depending on whether 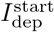 is smaller or larger than the corresponding value in the original model, respectively (see colored curves and dots in Fig. 6).

The above results show that all the spike times can be modulated in a predictable way by choosing different values of 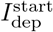 and/or 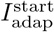, provided that condition (15) is satisfied.

#### 3.1.2 Temporal shift of all spike times except the first one

In this section, we examine mathematical procedures allowing to delay/anticipate all spike times except the first one. This can be obtained in two ways: either modifying 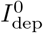 or modifying the parameters *c* and *η* of the Monod function (16).

In the first case, considering Eq. (26) and fixing the initial values of *V*^0^ and 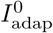, we have that increasing values of 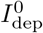 result in decreasing *ISI*s, and vice versa (see Figs. 7). We remark that shifting the first spike time automatically induces a shift of all the subsequent spike times. This in turn modifies the values of 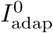 given by the Monod function (16); such effect, however, mainly depends on the monotonicity properties of the Monod function and of the potential *V* with respect to 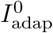. In the second case, recalling Eq. (16), we have

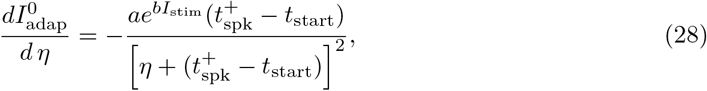

i.e. the function 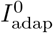 is decreasing or increasing with respect to *η* when *a* > 0 or *a* < 0, respectively. Consequently, if we fix the initial values of *V*^0^ and 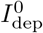 and the parameters *a*, *b*, *c*, increasing values of *η* will result in increasing *ISI*s when *a* < 0 and in decreasing *ISI*s when *a* > 0 (see Supp. Fig. 3). It should be noted that the Monod functions (16) obtained by modifying the parameter *η* have the same plateau value *c* + *ae*^*bI*_stim_^ (see Supp. Fig. 4); this implies that condition (15) is automatically satisfied.

**Figure 7:**
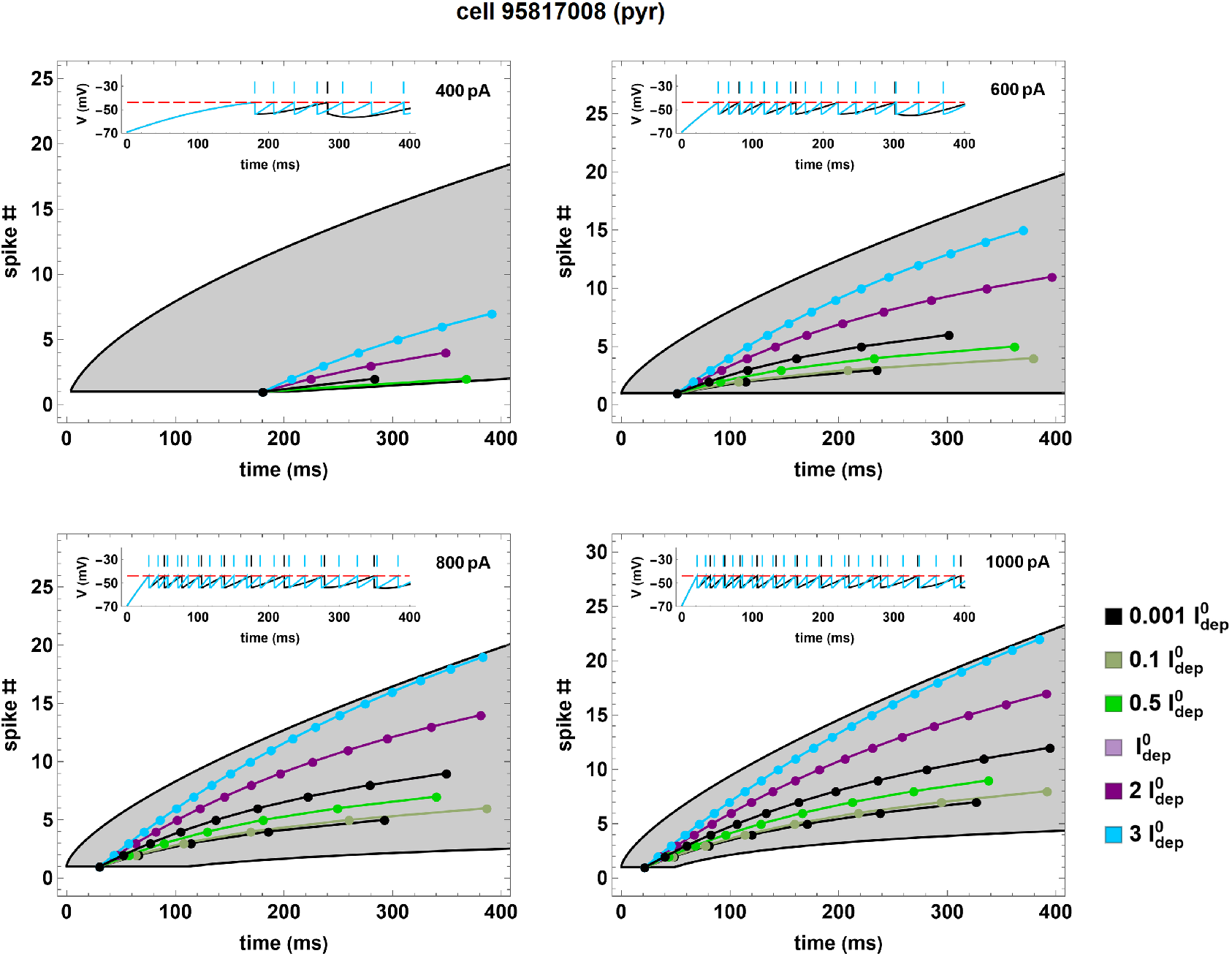
Spike number as a function of time for the pyramidal cell 95817008 (black curves and dots) and for 5 neuron copies (colored curves and dots) obtained by fixing the numerical values of all parameters except 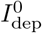. For all copies condition (15) is verified. The inset in each panel represents the original simulation compared with the copy corresponding to the 3 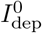.

In contrast, a change in the *c* parameter results in a translation of the Monod function along the *y*–axis, and therefore in a different value of the plateau (see Supp. Fig. 5). However, in this case to satisfy condition (15) for all injected currents *I*_stim_ it is sufficient to perturb *c* as follows

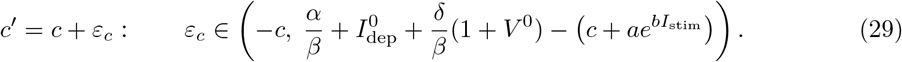

In Supp. Fig. 6 we show an example illustrating the effect of this perturbation.

#### 3.1.3 Changing the number of spikes

The goal of this section is to define mathematical procedures leading to an increase/decrease in the total number of spikes. This can be achieved either by modifying the time interval in which the Monod function (16) is defined (i.e. when a firing block occurs in response to a stimulation current *I*_stim_) or by varying the nondimensional parameter *α* in Eq. (7).

The first procedure is realized by modifying the coefficients *A_I,II_*, *B_I,II_* in (19). In particular, by introducing in (19) the perturbed parameters

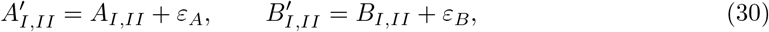

we obtain a firing block in the stimulation interval [*t*_start_, *T*] and in response to the stimulation current *I*_stim_ at the time

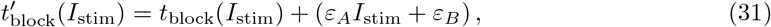

where *t*_block_ is the time in which the firing block occurs for the original model in response to the stimulation current *I*_stim_, i.e.

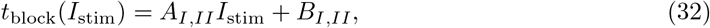

if and only if *ε_A_*, *ε_B_* are chosen such that

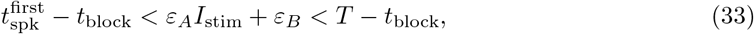

where 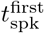 is the time of the first spike in response to the stimulation current *I*_stim_.

In particular, we have

- 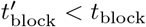 when 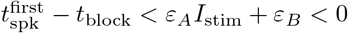;
- 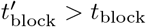 when 0 < *ε_A_I*_stim_ + *ε_B_* < *T* – *t*_block_.

We note that, with this approach, the range of currents in which the firing block occurs remains unchanged. However, in the copy generation procedure it is also possible to eliminate one or both firing block effects or to modify the value of 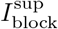 and 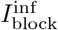 in Eq. 21 (see Fig. 8). The perturbation of each type of block (superior or inferior) is characterized by the triplet 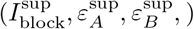 or 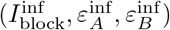, respectively. In particular, since the spike times are not affected by such perturbation, plots as the ones shown in Fig. 8 (consisting in variations of Input/Output (I/O) curves) help us emphasizing how the firing block perturbations illustrated above affect the number of spikes for different stimulation currents. An analogous variation in the I/O behaviour of a given neuron is shown in Fig. 9. There, we present the results obtained by changing, one parameter at the time, the overall shape of the excitability curve in such a way to: 1) cover the entire experimental variability range (shaded area in Fig. 9), starting from the optimization of a given experimental neuron (Fig. 9, black in top panel), 2) invert the I/O response (Fig. 9, middle panel), or 3) change the I/O curvature (Fig. 9, bottom panel). Similar results (see Suppl. Fig. 7) can be obtained using several other parameters’ perturbation.

**Figure 8:**
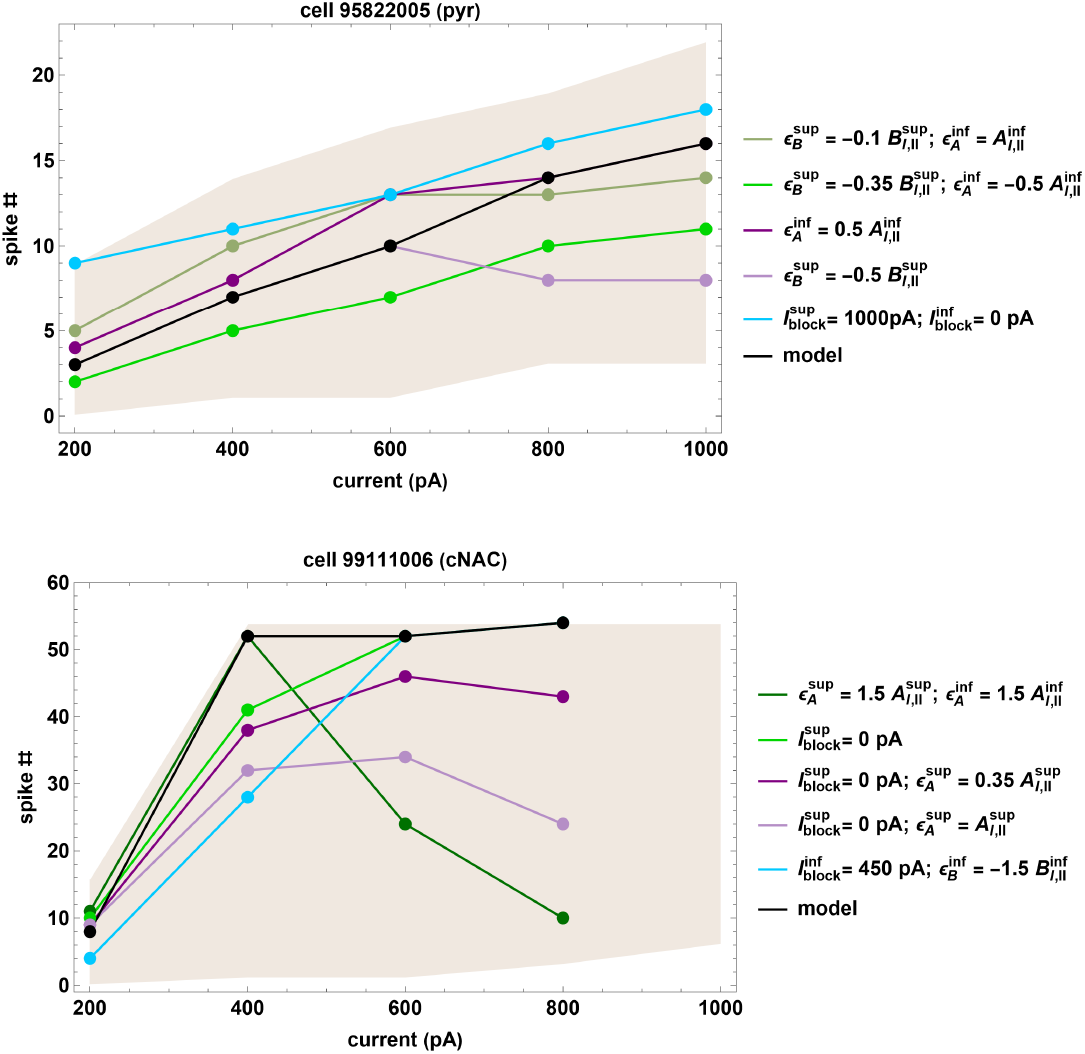
Effects of firing block perturbation. The two panels show the result of obtained using different combinations of firing block parameters, starting from two original optimizations (cell 95822005 and 99111006).

**Figure 9:**
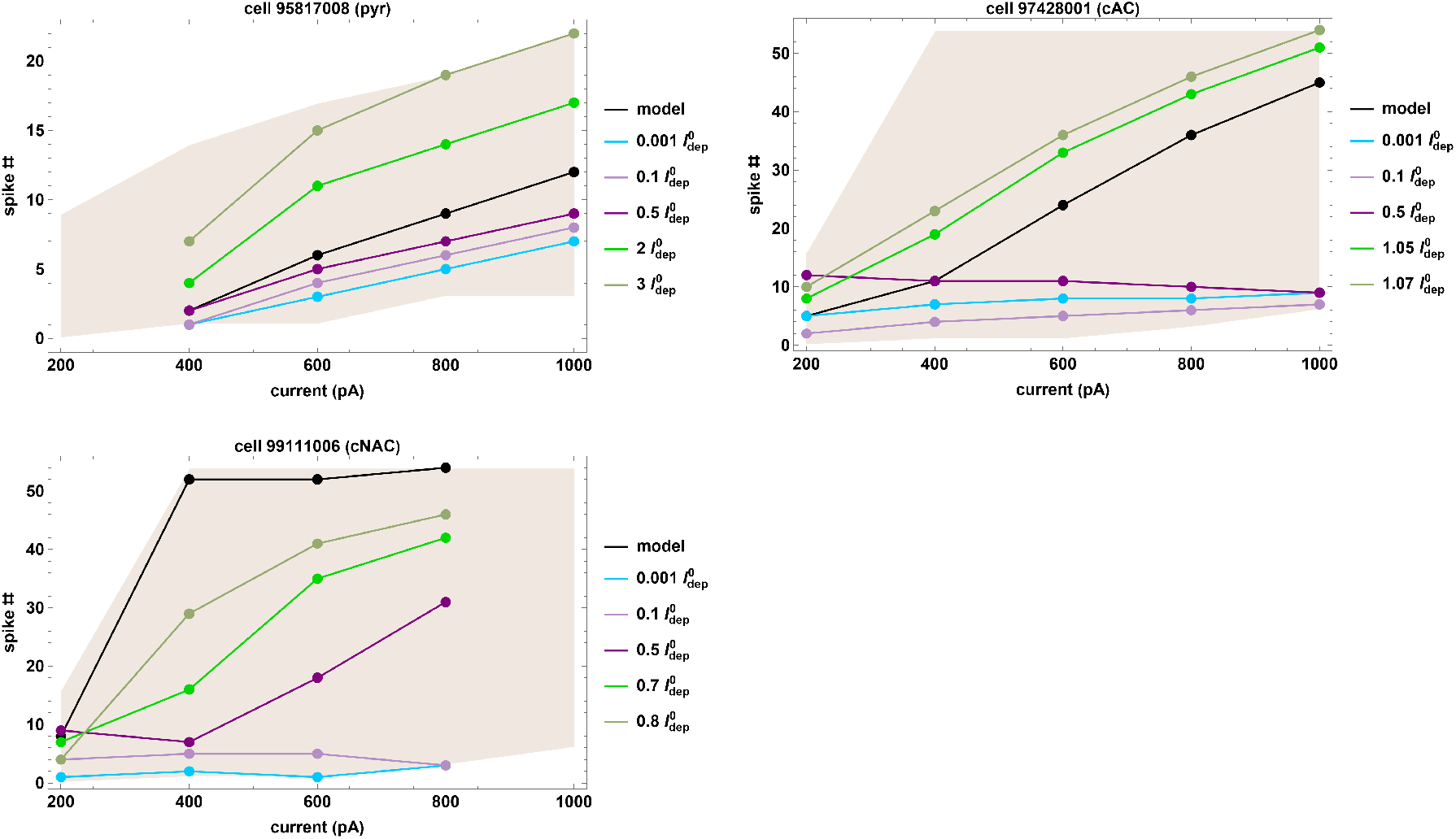
Number of spikes as a function of the input current. Number of spikes as a function of different constant currents for a pyramidal neuron (top left, black curves and dots), a cAC interneuron (top right, black curves and dots), and a cNAC interneuron (bottom, black curves and dots). Data related to copies obtained by fixing all parameters except for 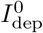 are shown via colored curves and dots.

In Figure 11 we compare the spike number as a function of the spike times for the A-GLIF model of the pyramidal cell 95817008 with those obtained for each of the 90 neuron copies. Note the large variability, in response to the same input, observed for the neuron copies.

**Figure 10:**
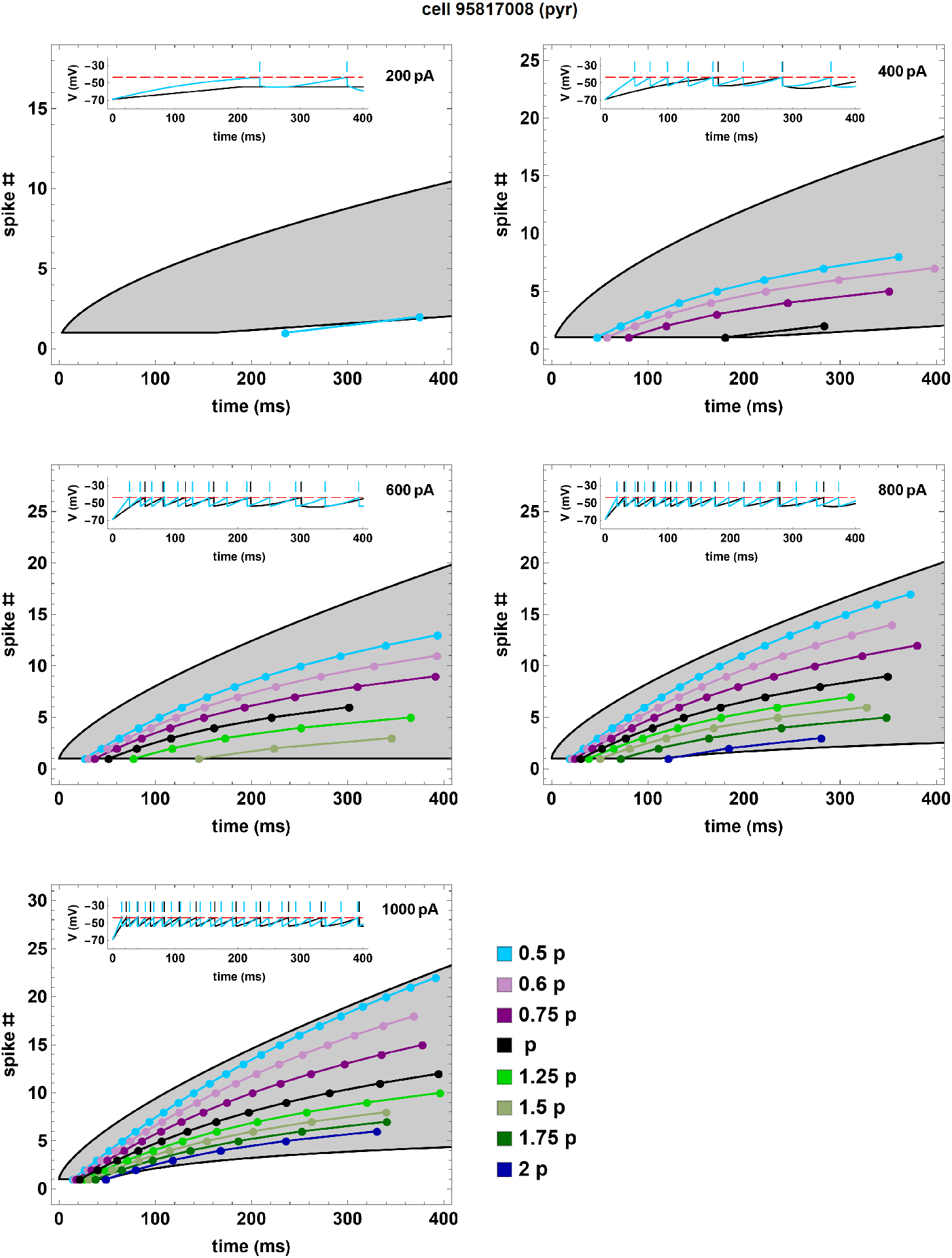
Spike number as a function of time for the pyramidal cell 95817008 (black curves and dots) and for 7 neuron copies (colored curves and dots) obtained by fixing all parameters except for *α*. The inset in each panel represents the original simulation compared with the copy corresponding to 0.5 *p*.

**Figure 11:**
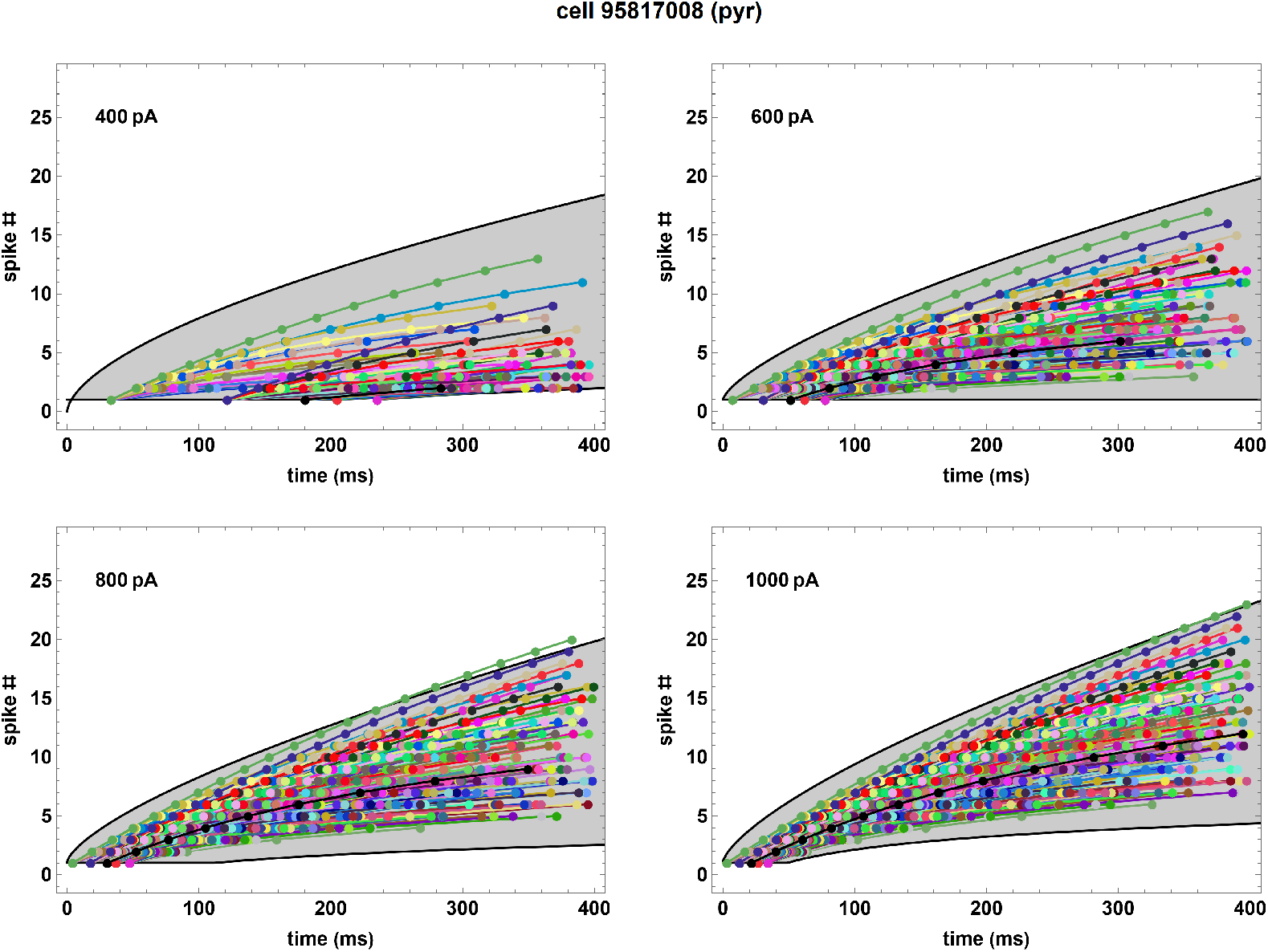
Spike number as a function of the spike times for the pyramidal cell 95817008 (black curves and dots) and for 90 neuron copies (colored curves and dots) obtained from it by fixing the numerical values of 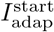 and *c* and varying the other parameters as follows: *h* 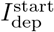 where *h* = 0.5, 0.75, 1, 2, 10; *m* 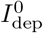 where *m* = 0.1, 0.25, 0.5, 1, 1.5, 2; *n η* where *n* = 0.75, 1, 2. The original model is obtained by setting *h* = *m* = *n* = 1 (black curves and dots).

Another way to change the total number of spikes is modifying the nondimensional parameter *α* while keeping *β*, *δ*, and *τ* constant (see Equation (7); we recall that *K* is a positive scaling constant defined as *K* = –*C_m_ E_L_ k*_2_). The variations on *α* is therefore going to be interpreted as variations in the dimensional parameters *C_m_* (whose values are derived from optimization procedures), whereas *Ṽ*_th_ and *E_L_* are considered constant due to their biophysical meaning. We use the superscript ^*n*^ and ^*o*^ to indicate the “*new*” and “*old*” value (for the neuron copies) of the corresponding parameter, respectively.

In particular, we consider 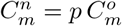 with *p* ≠ 1 being a positive parameter, whereas *k*_2_, *k*_1_, and *τ_m_* maintain the same value. This implies 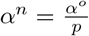, while *β*, *δ*, and *τ* remain unchanged. Consequently, the stability conditions (13) are identically satisfied. An important consequence of considering the updated parameter 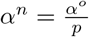 is that the threshold of the stimulation current *I*_th_ must also change accordingly by introducing 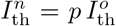. We provide here an example for sake of clarity. Let us consider a neuron satisfying 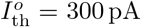; this implies that when the neuron is stimulated with a current *I*_stim_ = 200 pA it will not fire, whereas for *I*_stim_ = 400 pA the number of spikes will be non-zero – e.g., let us suppose it will be equal to 4. If we create a copy of such neuron with *p* =1/2 (while leaving the other parameters and initial data unaffected), we have that 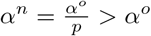. Since the potential *V* is increasing with respect to *α* (see Section 2.2.4), we have that the neuron copy will produce more spikes than the original one. In particular, when *I*_stim_ = 200 pA, the copy will behave as the original neuron behaved for 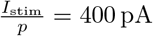, i.e. will produce 4 spikes. Therefore, the new threshold stimulation current cannot be still considered to be equal to 300 pA, and will have to be decreased proportionally to *p* as 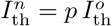 – which is equal to 150 pA. This assumption on 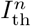 ensures that the condition in Equation (14) is automatically satisfied for the neuron copy both for any *p*, since 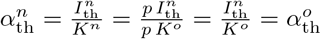.

We can therefore summarise the effects produced by considering neuron copies with *p* < 1 and *p* > 1 as follows (see Fig. 10 for an example):

1. for *p* < 1 we have *α^n^* > *α^o^* and 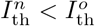. We hence have that the new initial condition on the first interval (defined in Equation (2)) 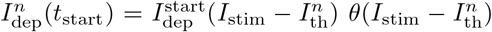 is greater than 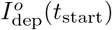. The fact that the potential *V* is a monotonically increasing function w.r.t. both 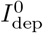 and *α* (see Section 2.2.4) implies that neuron copies constructed with *p* < 1 always exhibit more spikes than the original neuron.
2. for *p* > 1 we have *α^n^* < *α^o^* and 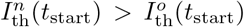. Therefore, we obtain that here 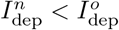. The abovementioned monotonicity properties of *V* w.r.t. both 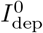 and *α* imply that, in this case, neuron copies constructed with *p* > 1 always exhibit less spikes than the original neuron.

Finally, we note that the parameter *τ* also affects the spiking dynamics, but only on the nondimensional level, since it only influences the nondimensional time scalings (see Supplementary Materials).

The overall conceptual results of Sections 3.1.1–4 are summarized in Table 2, where we report a qualitative description of the effects obtained by perturbing different parameters. Additionally, Fig. 11 and Supp. Fig. 8 provide 90 and 108 neuron copies, respectively, obtained by implementing all parameter variations described in Table 2.

**Table 2:** Summary of the possible variations in the copies firing dynamics linked to the mathematical procedure for each effect as illustrated in Sections 3.1.1–4.

| Effect | Parameter variation |
| --- | --- |
| **Temporal shift of all spike times** |  |
| Delay the first spike time and changes all other spike times | Increase $I_{\text{adap}}^{\text{start}}$ |
| Anticipate the first spike time and changes all other spike times | Increase $I_{\text{dep}}^{\text{start}}$ |
| **Temporal shift of all spike times except the first one** |  |
| Anticipate all but the first spike time | Increase $I_{\text{dep}}^0$ |
| Delay all but the first spike time | Increase $c$ |
| Delay/anticipate all but the first spike time if $a > 0$ or $a < 0$, respectively | Increase $\eta$ |
| **Change in number of spikes** |  |
| Generate more/less spikes | Positive/negative values of $\varepsilon_A I_{\text{stim}} + \varepsilon_B$ for any $I_{\text{stim}}$ |
| Increase the total number of spikes | Increase $\alpha$<br>(⇒ decrease $I_{\text{th}}$) |

### 3.2 Numerical implementation of the generation procedure

The aim of this section is to illustrate the numerical procedure leading to the generation of neuron copies, and thus a downloadable database, as summarized in Fig. 12.

**Figure 12:**
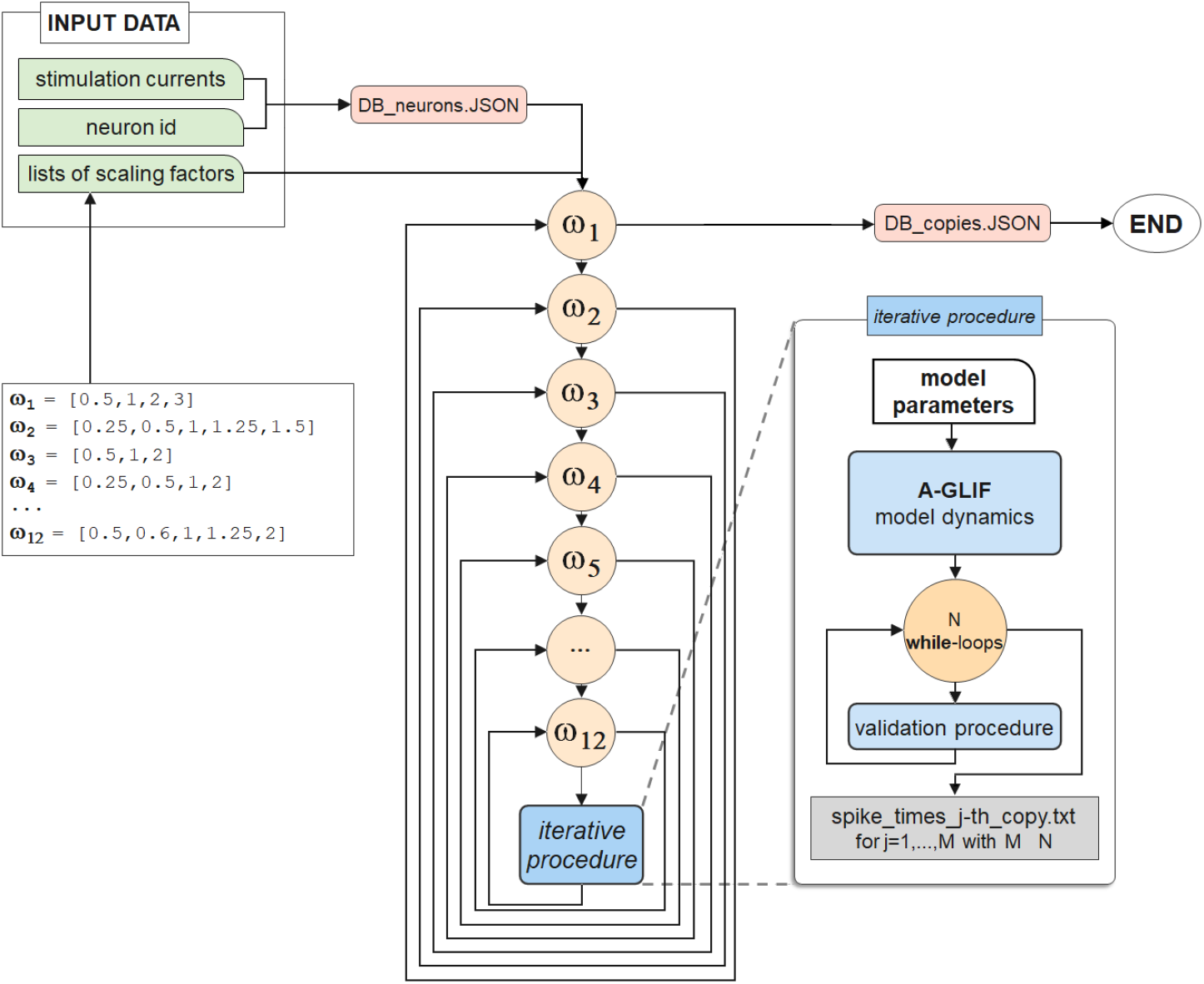
Flow-chart illustration of the numerical implementation for the generation of a database of neuron copies. Here the sets *ω_i_*, *i* = 1,…, 12 represent the lists of scaling factors leading to modified values of the corresponding parameter Γ_*i*_, representing a generic parameter among the ones in Eq. (34). The initial file `DB_neurons_JSON` containing information about experimental and optimized pyramidal neurons and interneurons is then fed into 12 nested for-loops and, after an A-GLIF simulation and a validation procedure, leads to a new file `DB_copies_JSON` containing information about parameter values and simulated spike times for the generated *M* admissible copies.

In line with the investigation carried out in Section 3.1, copies of pyramidal neurons and interneurons are generated starting from the optimized values of the following 12 parameters^2^

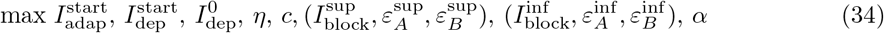

via an iterative procedure, where the input value of each parameter is altered by considering a new, rescaled value based on an admissible scaling factor contained in a suitable list of variable size.

A neuron copy is considered admissible if and only if the resulting spike times at each constant stimulation current fall within the corresponding experimental region as represented in Fig. 1 (see Supporting information). Therefore, a validation procedure is run during the copies generation in order to dismiss those for which this requirement does not hold. In this procedure, the set of parameters used for the copy generation is discarded as soon as the numerical spike times, for one or more stimulation currents, lie outside the region defined by experimental data.

In the case of parameters max 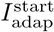 and 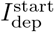 it is possible to perform an *a priori* validation study based on the experimental range of first spike time at each current in order to identify ranges of variability for the corresponding scaling factors. This procedure (described into the Supplementary information) allows to reduce the number of dismissed copies resulting from the validation procedure described above, that must be executed nevertheless.

A schematic representation of the implementation procedure is outlined in Fig. 12. Experimental data^3^ and optimized parameter values for 84 pyramidal neurons and interneurons (identified by a neuron ID) are stored into a `JSON` file in a database like structure designed to efficiently retrieve data necessary to calculation. For each parameter in Eq. (34) here generically defined as Γ_*i*_, *i* = 1,…, 12, suitable lists *ω_i_* of scaling factors *h_i,j_* are defined as

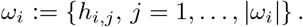

The validation procedure described above will allow to select the admissible coefficients *h_i,j_*, i.e. the scaling factors for which the new, rescaled parameter *h_i,j_*Γ_*i*_ satisfies the admissibility constraints. We note that for *h*_1,*j*_ = 0, and *h_i,j_* = 1 for *i* ≠ 1 we retrieve the value of the parameter Γ_*i*_ corresponding to the reference neuron from which the copies are derived. The number of copies generated depends on the number of admissible scaling factors considered for each parameter, up to a maximum of 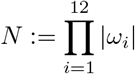. Within these 12 nested for-loops, the A-GLIF simulations and the validation procedure described above are performed: this finally leads to set the parameters for *M* ≤ *N* admissible copies which are then saved together with simulated spiking times into an analogous `JSON` data structure.

Figure 13 shows a snapshot of the user interface for entering the required information (neuron type and the number of neuron copies) and to visualize the resultant plots of the spike number as a function of the time for the neuron copies. All generated data can be downloaded as a `JSON` file.

**Figure 13:**
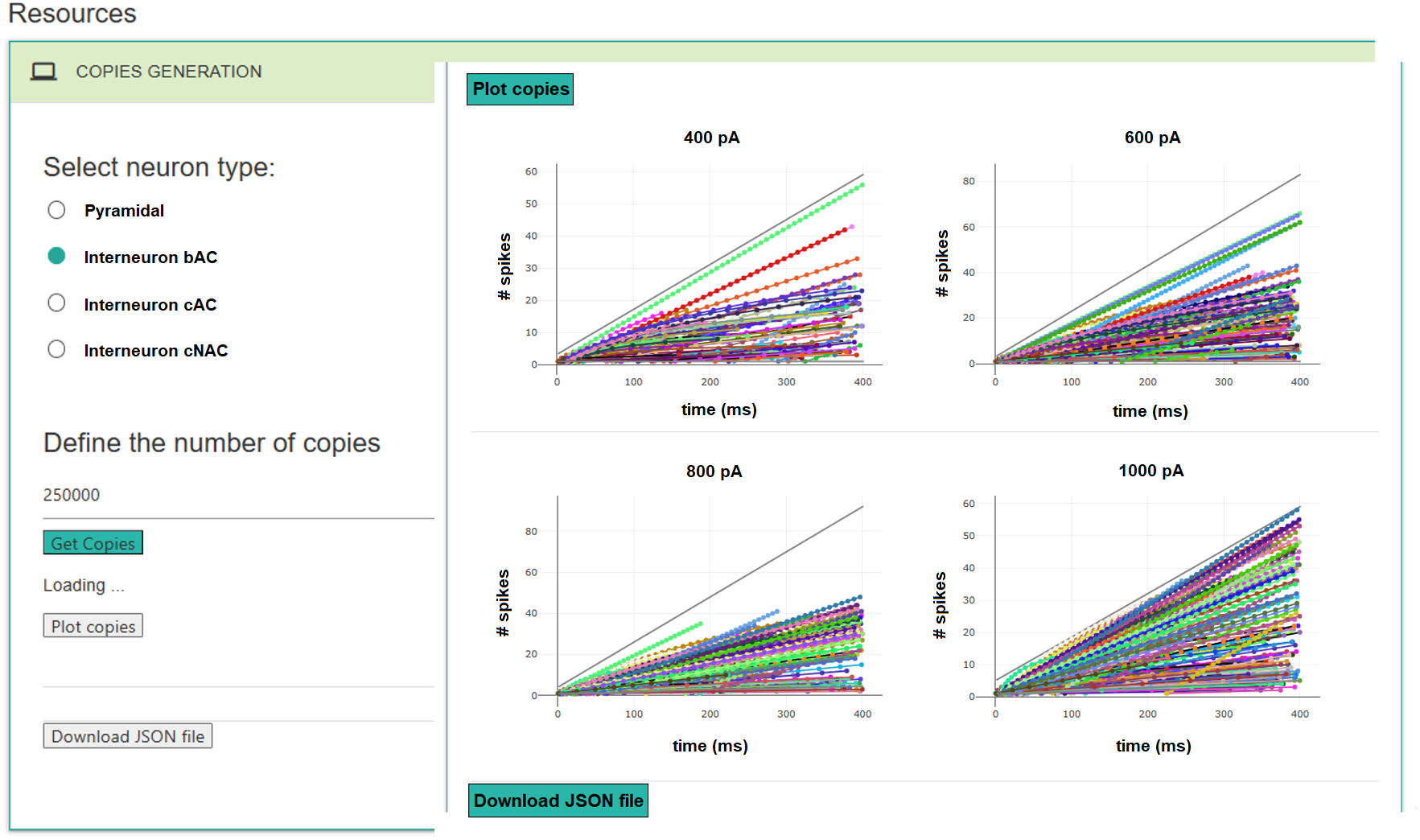
Copies generation frontend. The user can select the neuron type and set the number of neuron copies. By clicking on *Plot copies* and *Download JSON file* the user can visualise the plots of the spike number as a function of the time for the neuron copies and download all data saved in JSON format, respectively.

## 4 Discussion

In this work, we have introduced an automatic numerical procedure to generate neuron copies based on experimental and optimized data for CA1 pyramidal neurons and interneurons. Our method relies on the simulation of neuron firing dynamics by means of the A-GLIF modelling framework, because of its ability to effectively capture such dynamics (and being computationally efficient), and of its analytical properties (which allow to control the copies firing behaviour).

The neuron copies can be obtained by varying the 12 parameters of any given optimized model via scaling factors which, through a validation procedure, ensure that their firing properties remain within the observed experimental ranges. The parameters which can be potentially modified here are the initial conditions relative to the A-GLIF Cauchy problems (3 parameters), the Monod parameters defining the update value of the adaptation current (2 parameters), the firing blocks parameters determining the time interval in which the Monod function should be defined (6 parameters), and the internal nondimensional parameter *α*. These alterations lead to changes in both the spike times and the number of spikes, as summarized in Table 2.

The generation procedure introduced here allows to obtain an arbitrarily large number of heterogeneous neuron copies with a predictable dynamical behaviour and falling within the experimental ranges of CA1 pyramidal neurons and interneurons. We provide approximately 500K predetermined copies (including the computed spike times) in a JSON database which can be downloaded from the *live paper section of EBRAINS* (see Fig. 13).

Further experimental data, also related to other types of neurons, will allow us, in the future, to broaden such database and extend our approach to construct neural networks for different brain areas as well.

## Supporting information

**Supplementary Figure 1:**
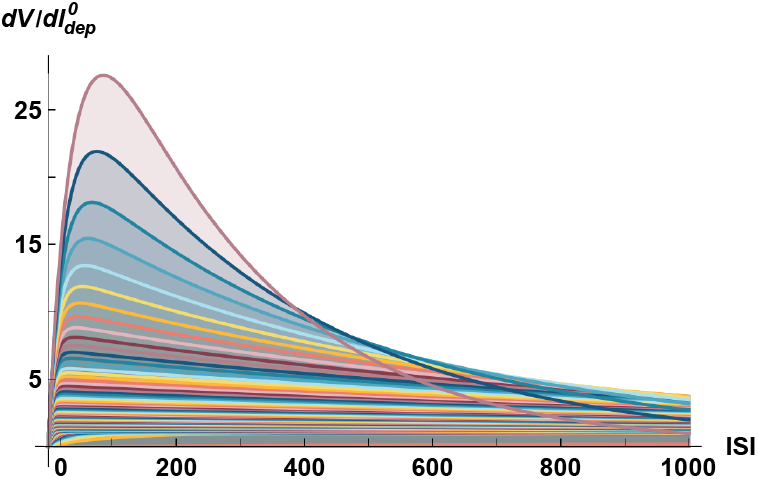
Plot of the derivative of *V* with respect to 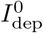. Plot of the derivative (26) as function of *t* – *t*_0_ obtained imposing the conditions (13) with a step of 0.01 for both parameters *β* and *δ*.

**Supplementary Figure 2:**
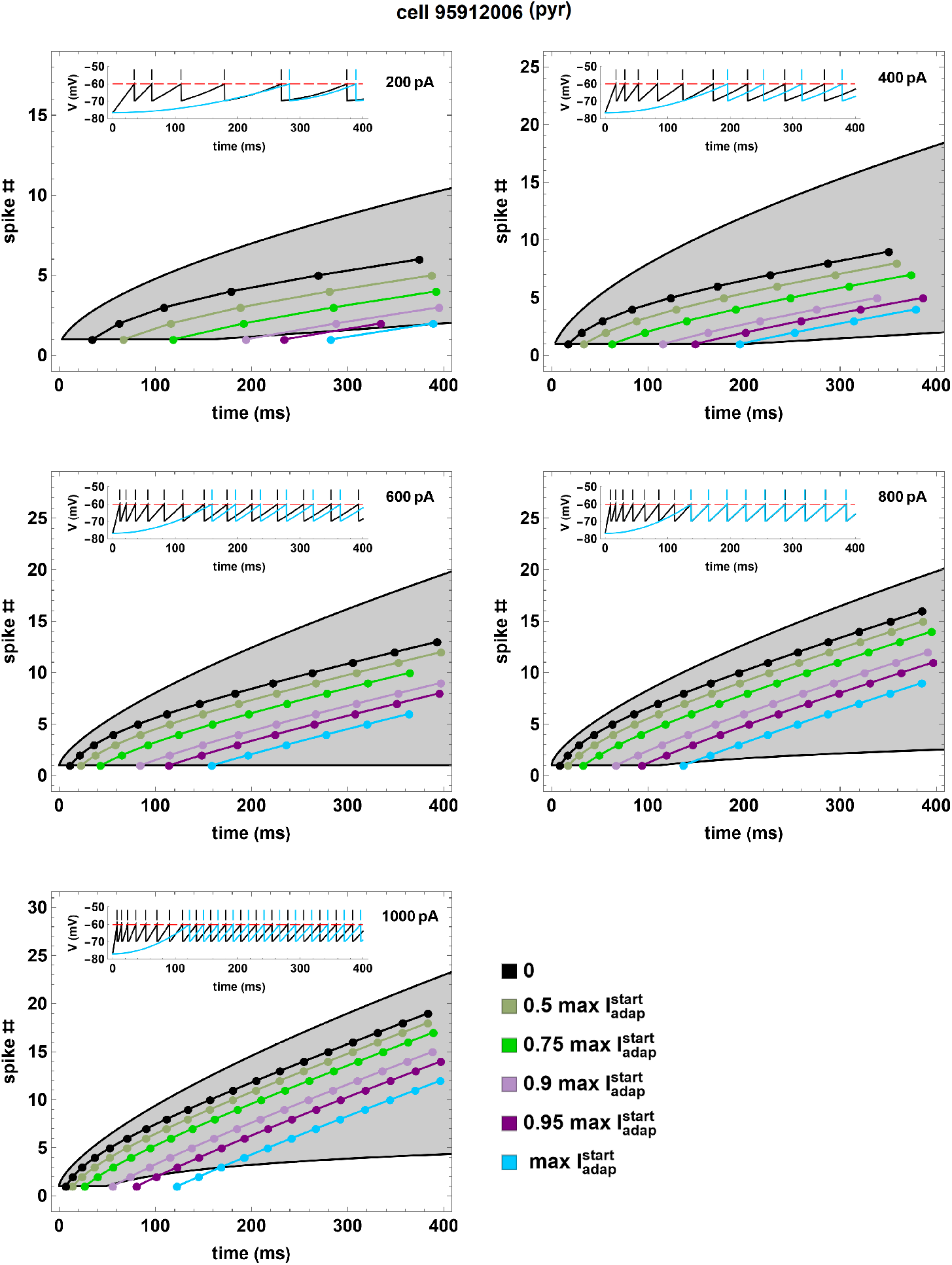
Spike number as function of time for the pyramidal cell 95912006 on varying 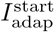. Spike number as a function of time for the pyramidal cell 95912006 (black curves and dots) and for 5 neuron copies (colored curves and dots) obtained by fixing the numerical values of all parameters except 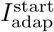. For each current *I*_stim_ we set max 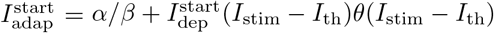. The gray areas cover the experimental regions of the data. The inset in each panel represents the original simulation of the firing dynamics compared with the copy corresponding to the max 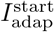.

**Supplementary Figure 3:**
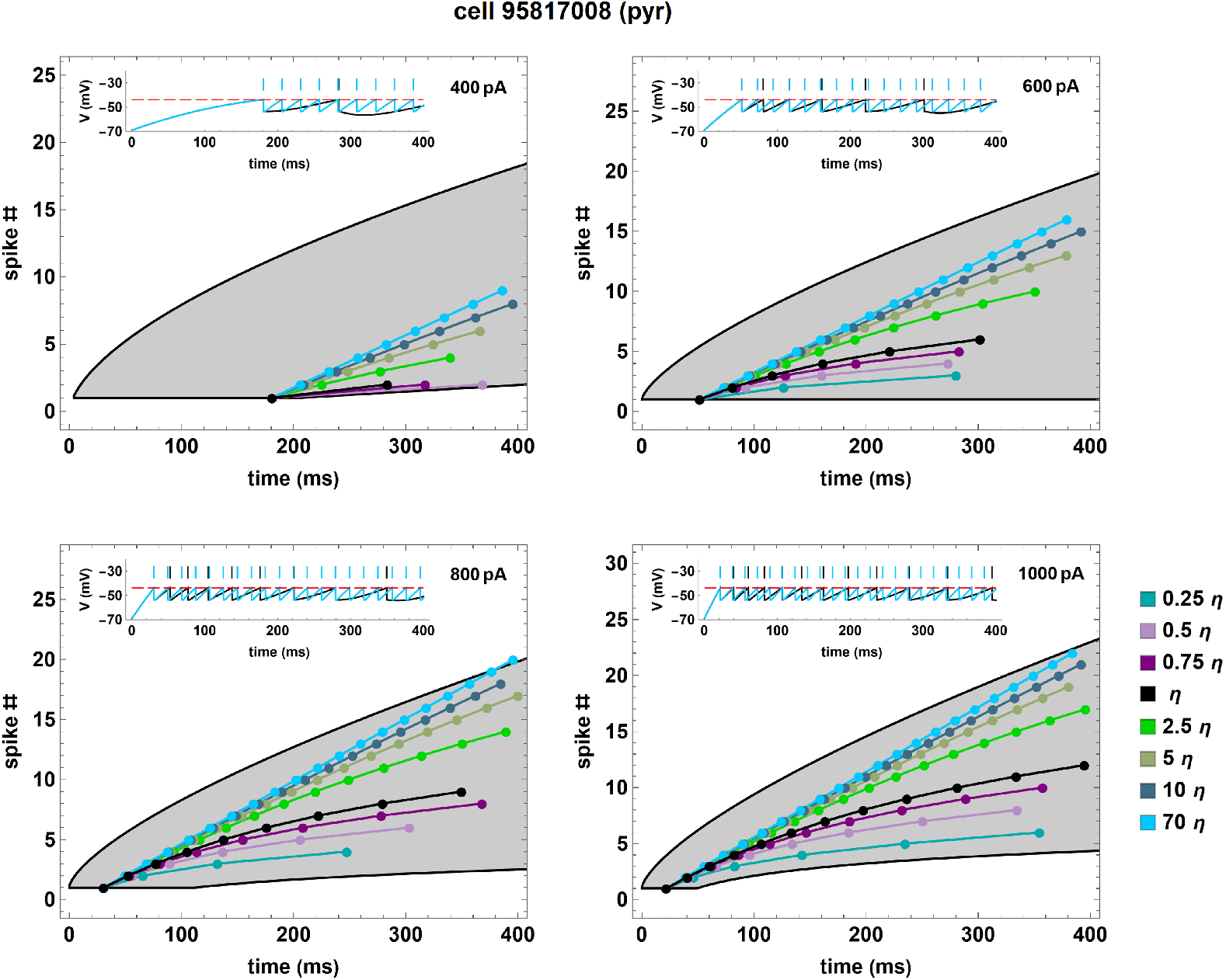
Spike number as function of time for the pyramidal cell 95817008 on varying *η*. Spike number as a function of time for the pyramidal cell 95817008 (black curves and dots) and for 7 neuron copies (colored curves and dots) obtained by fixing all parameters except *η* in the Monod function (16). The inset in each panel represents the original simulation compared with the copy corresponding to 70 *η*.

**Supplementary Figure 4:**
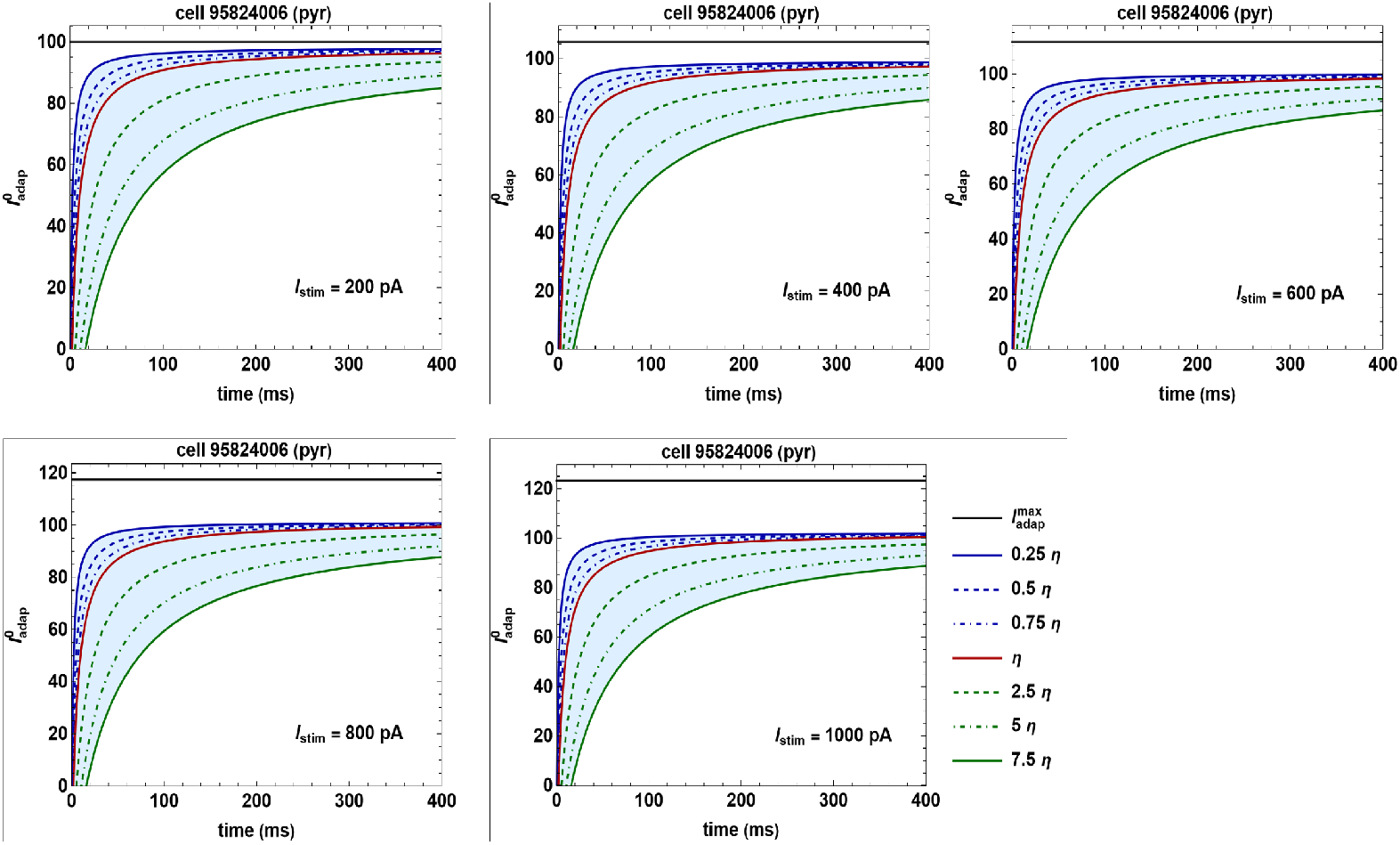
Plot of the Monod functions for the pyramidal cell 95824006 on varying the parameter *η*. Plots of the Monod function (16) for the pyramidal cell 95824006 (red) and for the perturbed functions (blue and green) obtained on varying the parameter *η*. For each *I*_stim_ we set 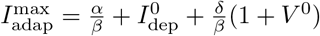. In all cases *a* > 0.

**Supplementary Figure 5:**
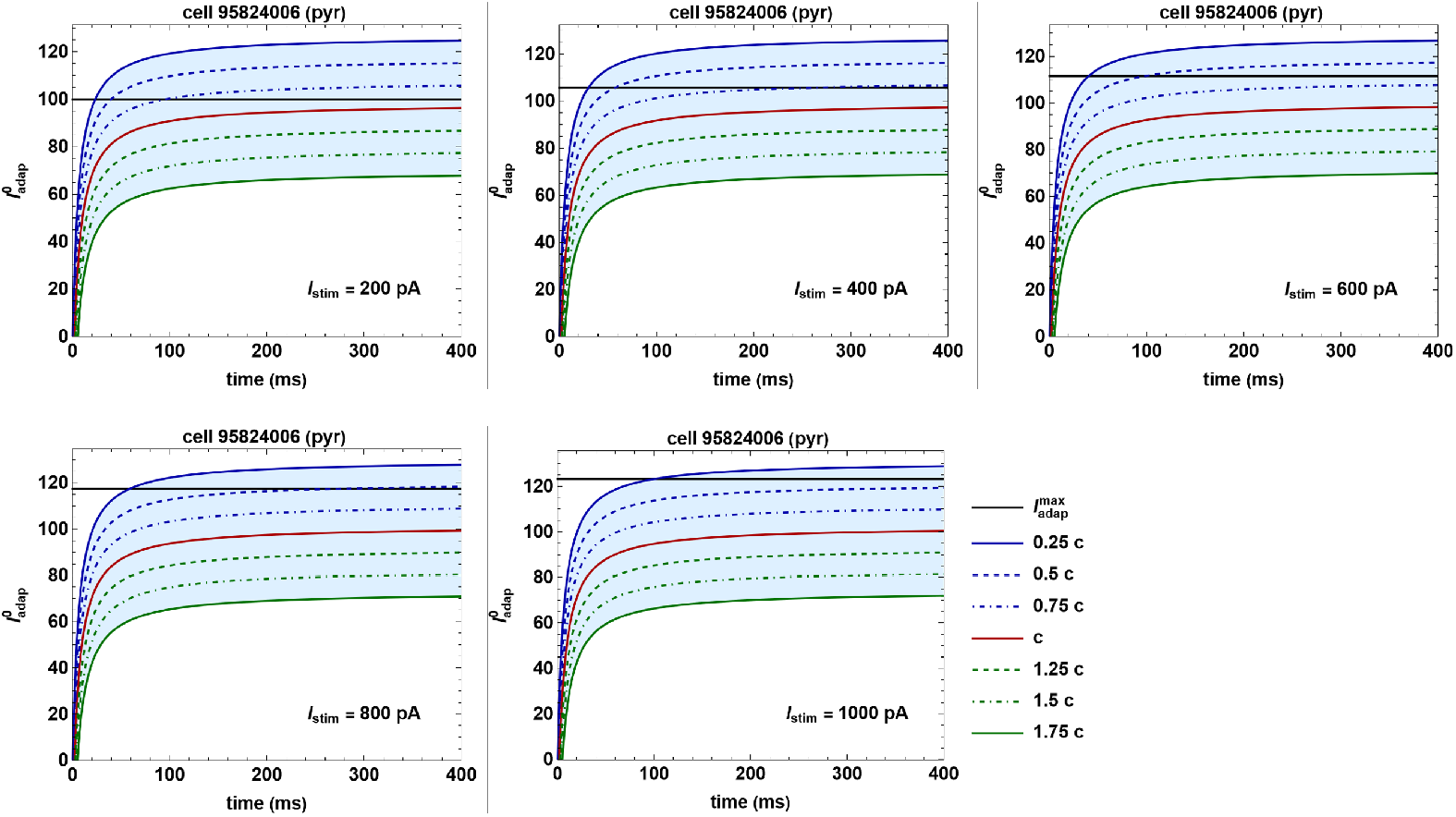
Plot of the Monod functions for the pyramidal cell 95824006 on varying the parameter *c*. Plots of the Monod function (16) for the pyramidal cell 95824006 (red) and for the perturbed functions (blue and green) obtained on varying the parameter *c*. For each *I*_stim_ we set 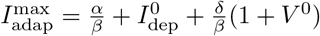. In all cases *a* > 0.

**Supplementary Figure 6:**
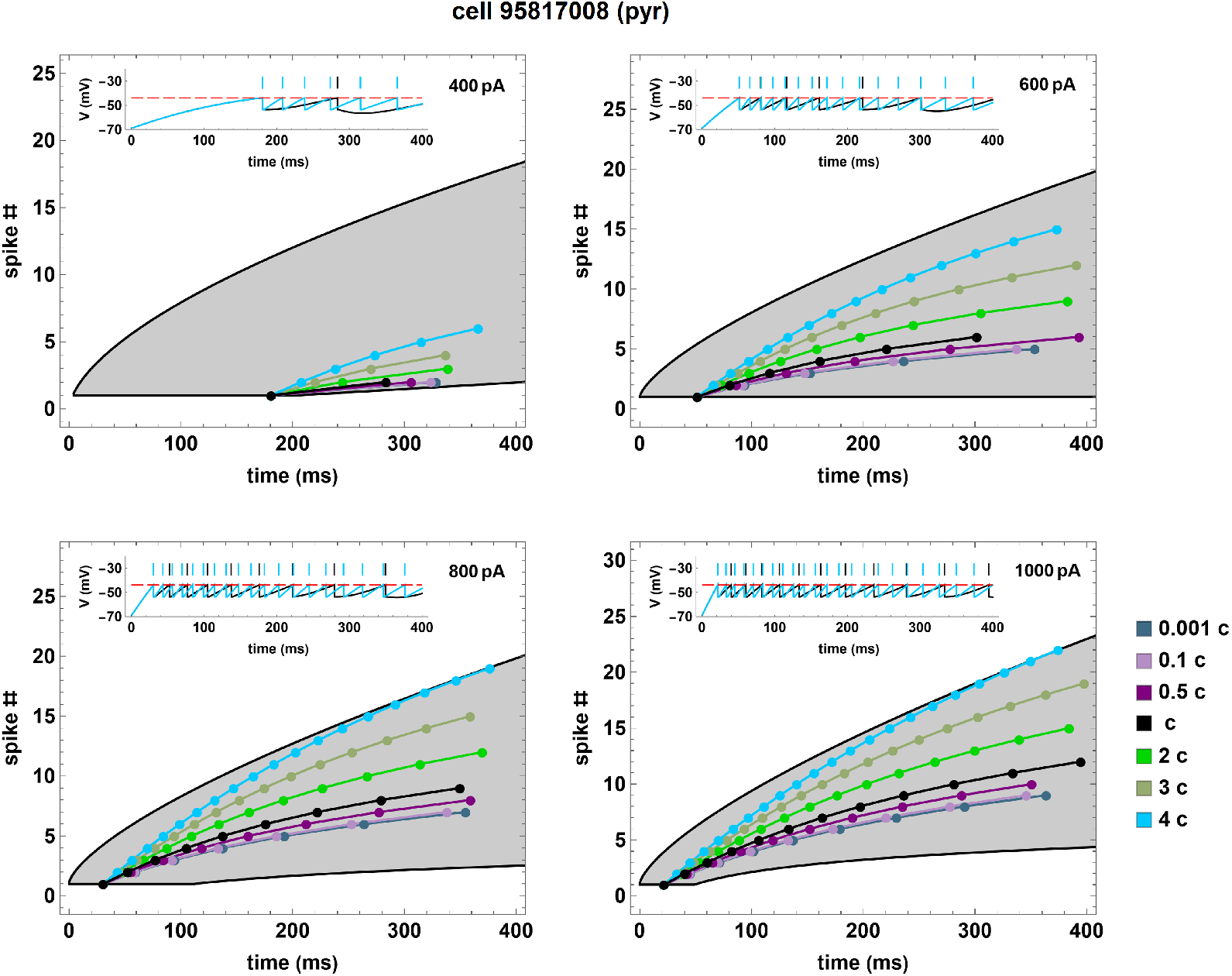
Spike number as function of time for the pyramidal cell 95817008 on varying *c*. Spike number as a function of time for the pyramidal cell 95817008 (black curves and dots) and for 6 neuron copies (colored curves and dots) obtained by fixing all parameters except for *c* in the Monod function (16). The inset in each panel represents the original simulation compared with the copy corresponding to 4 *c*.

**Supplementary Figure 7:**
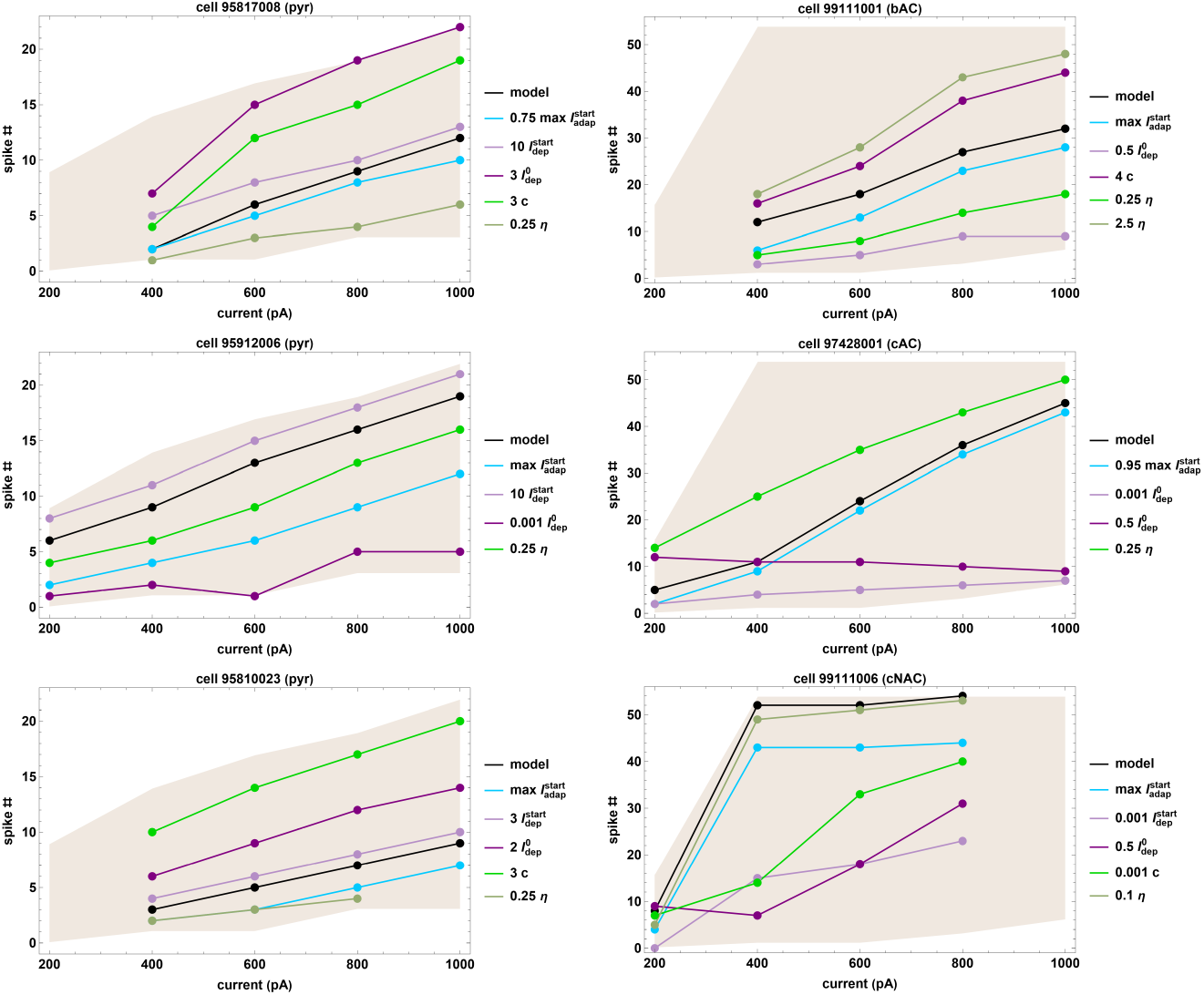
Number of spikes as function of the constant stimulation currents. Number of spikes as function of different constant currents (black curves and dots) for three pyramidal neurons (left column) and three interneurons of bAC, cAC and cNAC type (right column, top, middle, and bottom, respectively) and for some copies (colored curves and dots) obtained by fixing the numerical values of all parameters except the ones indicated in the legends.

**Supplementary Figure 8:**
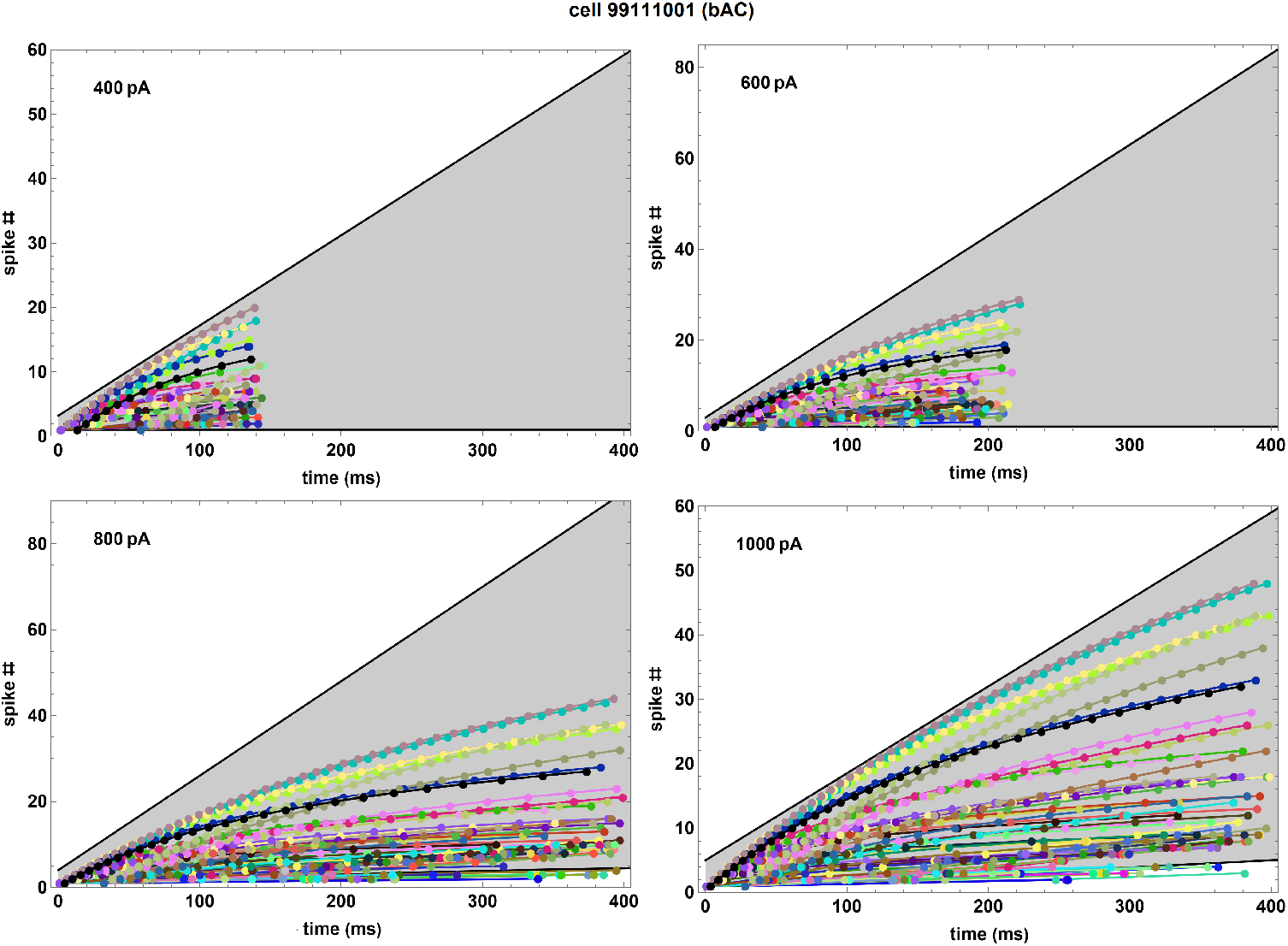
Spike number as function of time for the interneuron 99111001 on varying different parameters. Spike number as a function of the spike times for the interneuron 99111001 (black curves and dots) and for 108 neuron copies (colored curves and dots) obtained from the former by varying the parameters as follows: *h* max 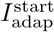 where *h* = 0, 1; *m* 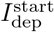, where *m* = 1, 10; *n* 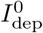 where *n* = 0.001, 0.5, 1; *p c* where *p* = 0.1, 1, 4; *q η* where *q* = 0.25, 1, 2.5. The original model is obtained by setting *h* = 0, *m* = *n* = *p* = *q* = 1 (black curves and dots).

### Temporal shift of all spikes at nondimensional level

We consider variations in the dimensional parameters *C_m_*, *k*_2_, *k*_1_, and *τ_m_* (whose values are again derived from optimization procedures) such that *α*, *β*, and *δ* remain constant. Moreover, we adopt the same notation of Section 3.1.3 regarding “*old*” and “*new*” parameter values.

In this case, we assume 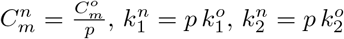, and 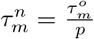, leading to 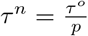 while *α*, *β*, and *δ* do not change. Therefore, in this scenario the threshold current *I*_th_ also remains constant. This implies that both the stability conditions (13) and the the conditions in Eq. (14) are automatically verified. In this scenario, the sole variation of the copies firing dynamics lies in the (nondimensional) *ISI*s as follows:

1. for *p* < 1 we have 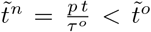, therefore the (nondimensional) temporal dynamics are shrinking;
2. for *p* > 1 we have 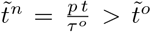, therefore the (nondimensional) temporal dynamics are dilating.

We remark that the changes on *τ* outlined above only affect the nondimensional time scalings; consequently, although they might lead in the case *p* < 1 to a reduced computational cost to reproduce the same firing patterns of the original neuron, they will leave the curves corresponding to dimensional firing dynamics unaffected.

### A priori validation for the perturbation of max 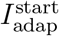 and 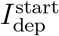

The procedure mentioned in Section 3.2 is based on pre-selecting at least 4 scaling factors for the corresponding parameter, according to which the first spike time is calculated. This allows us to construct an interpolating function by associating coefficient values to their corresponding time of first spike. Taking into account the experimental boundaries available for each stimulation current, it is possible to identify the range of admissibility values for the parameter scaling factors (see Supp. Fig. 9). These bounds are defined as piecewise curves

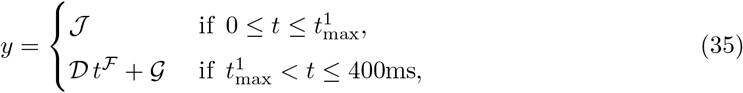

where *y* represents the number of spike, *t* the time, and 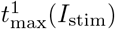 is the largest time supporting a single spike for the constant stimulation current *I*_stim_.

In particular, *y*_low_ (resp. *y*_up_) is the curve bounding the experimental region from below (resp. above) in the space of spike numbers vs time. The values of the coefficients 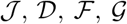, together with 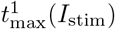, are provided in Table 1. The final interval of admissibility for max 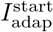 and 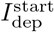 is then obtained by intersecting the intervals related to the different constant stimulation currents considered. We note that the admissibility ranges for max 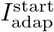 and 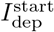 vary for each reference neuron we use for the copies generation procedure.

**Supplementary Table 1:**
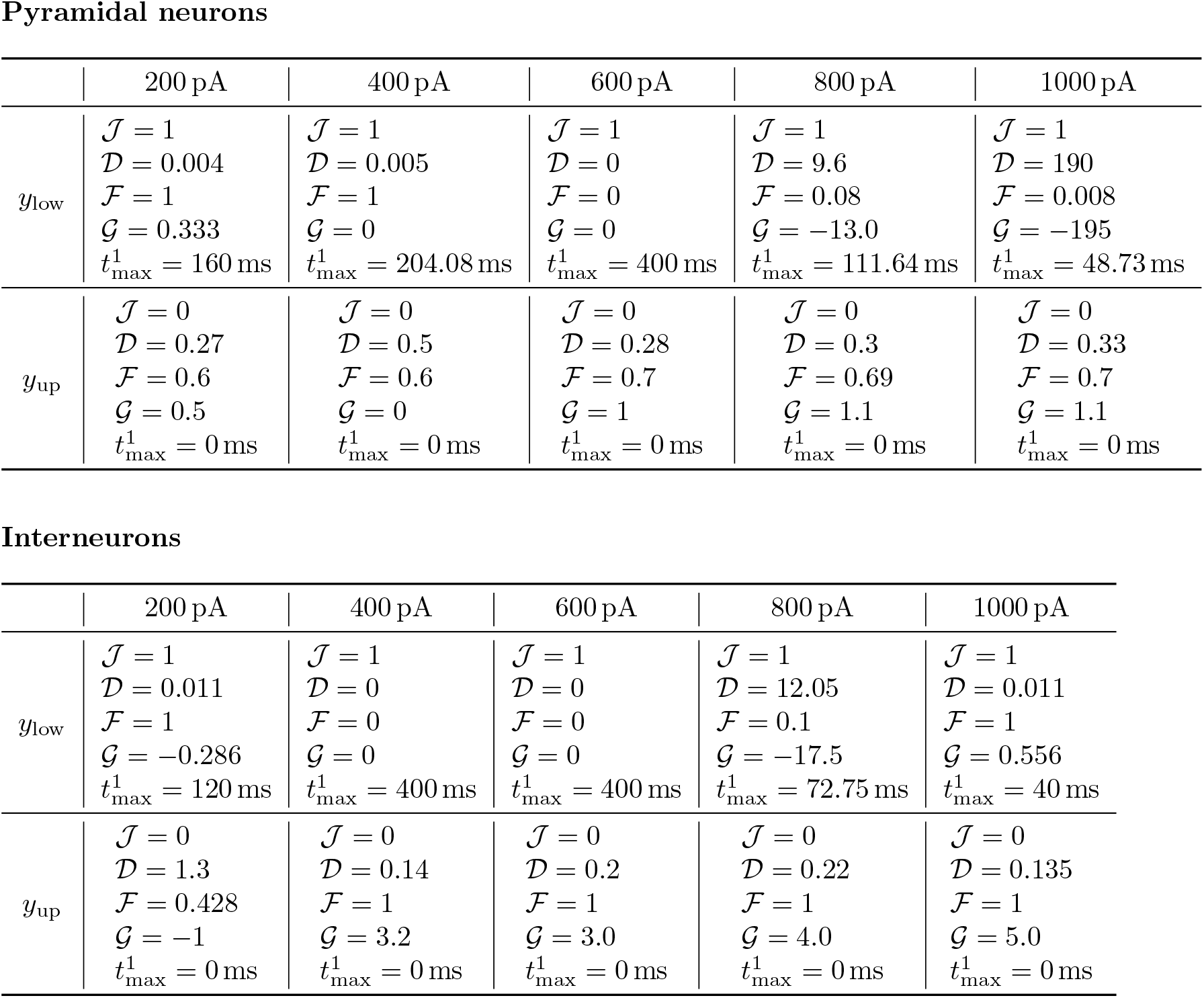
Coefficients of the lower and upper limit curves for the region of experimentally feasible spike number *y*_low_, *y*_up_, respectively, as function of time *t* defined in Eq. (35), together with the largest time supporting a single spike for each constant stimulation current *I*_stim_ = 200, 400, 600, 800, 1000 pA, both for pyramidal neurons (upper table) and interneurons (lower table).

Pyramidal neurons
|  | 200 pA | 400 pA | 600 pA | 800 pA | 1000 pA |
| --- | --- | --- | --- | --- | --- |
| $y_{\text{low}}$ | $\mathcal{J} = 1$<br>$\mathcal{D} = 0.004$<br>$\mathcal{F} = 1$<br>$\mathcal{G} = 0.333$<br>$t_{\text{max}}^1 = 160 \text{ ms}$ | $\mathcal{J} = 1$<br>$\mathcal{D} = 0.005$<br>$\mathcal{F} = 1$<br>$\mathcal{G} = 0$<br>$t_{\text{max}}^1 = 204.08 \text{ ms}$ | $\mathcal{J} = 1$<br>$\mathcal{D} = 0$<br>$\mathcal{F} = 0$<br>$\mathcal{G} = 0$<br>$t_{\text{max}}^1 = 400 \text{ ms}$ | $\mathcal{J} = 1$<br>$\mathcal{D} = 9.6$<br>$\mathcal{F} = 0.08$<br>$\mathcal{G} = -13.0$<br>$t_{\text{max}}^1 = 111.64 \text{ ms}$ | $\mathcal{J} = 1$<br>$\mathcal{D} = 190$<br>$\mathcal{F} = 0.008$<br>$\mathcal{G} = -195$<br>$t_{\text{max}}^1 = 48.73 \text{ ms}$ |
| $y_{\text{up}}$ | $\mathcal{J} = 0$<br>$\mathcal{D} = 0.27$<br>$\mathcal{F} = 0.6$<br>$\mathcal{G} = 0.5$<br>$t_{\text{max}}^1 = 0 \text{ ms}$ | $\mathcal{J} = 0$<br>$\mathcal{D} = 0.5$<br>$\mathcal{F} = 0.6$<br>$\mathcal{G} = 0$<br>$t_{\text{max}}^1 = 0 \text{ ms}$ | $\mathcal{J} = 0$<br>$\mathcal{D} = 0.28$<br>$\mathcal{F} = 0.7$<br>$\mathcal{G} = 1$<br>$t_{\text{max}}^1 = 0 \text{ ms}$ | $\mathcal{J} = 0$<br>$\mathcal{D} = 0.3$<br>$\mathcal{F} = 0.69$<br>$\mathcal{G} = 1.1$<br>$t_{\text{max}}^1 = 0 \text{ ms}$ | $\mathcal{J} = 0$<br>$\mathcal{D} = 0.33$<br>$\mathcal{F} = 0.7$<br>$\mathcal{G} = 1.1$<br>$t_{\text{max}}^1 = 0 \text{ ms}$ |

Supplementary Table 1: Coefficients of the lower and upper limit curves for the region of experimentally feasible spike number $y_{\text{low}}$ , $y_{\text{up}}$ , respectively, as function of time $t$ defined in Eq.
|  | 200 pA | 400 pA | 600 pA | 800 pA | 1000 pA |
| --- | --- | --- | --- | --- | --- |
| $y_{\text{low}}$ | $\mathcal{J} = 1$<br>$\mathcal{D} = 0.011$<br>$\mathcal{F} = 1$<br>$\mathcal{G} = -0.286$<br>$t_{\text{max}}^1 = 120 \text{ ms}$ | $\mathcal{J} = 1$<br>$\mathcal{D} = 0$<br>$\mathcal{F} = 0$<br>$\mathcal{G} = 0$<br>$t_{\text{max}}^1 = 400 \text{ ms}$ | $\mathcal{J} = 1$<br>$\mathcal{D} = 0$<br>$\mathcal{F} = 0$<br>$\mathcal{G} = 0$<br>$t_{\text{max}}^1 = 400 \text{ ms}$ | $\mathcal{J} = 1$<br>$\mathcal{D} = 12.05$<br>$\mathcal{F} = 0.1$<br>$\mathcal{G} = -17.5$<br>$t_{\text{max}}^1 = 72.75 \text{ ms}$ | $\mathcal{J} = 1$<br>$\mathcal{D} = 0.011$<br>$\mathcal{F} = 1$<br>$\mathcal{G} = 0.556$<br>$t_{\text{max}}^1 = 40 \text{ ms}$ |
| $y_{\text{up}}$ | $\mathcal{J} = 0$<br>$\mathcal{D} = 1.3$<br>$\mathcal{F} = 0.428$<br>$\mathcal{G} = -1$<br>$t_{\text{max}}^1 = 0 \text{ ms}$ | $\mathcal{J} = 0$<br>$\mathcal{D} = 0.14$<br>$\mathcal{F} = 1$<br>$\mathcal{G} = 3.2$<br>$t_{\text{max}}^1 = 0 \text{ ms}$ | $\mathcal{J} = 0$<br>$\mathcal{D} = 0.2$<br>$\mathcal{F} = 1$<br>$\mathcal{G} = 3.0$<br>$t_{\text{max}}^1 = 0 \text{ ms}$ | $\mathcal{J} = 0$<br>$\mathcal{D} = 0.22$<br>$\mathcal{F} = 1$<br>$\mathcal{G} = 4.0$<br>$t_{\text{max}}^1 = 0 \text{ ms}$ | $\mathcal{J} = 0$<br>$\mathcal{D} = 0.135$<br>$\mathcal{F} = 1$<br>$\mathcal{G} = 5.0$<br>$t_{\text{max}}^1 = 0 \text{ ms}$ |

**Supplementary Figure 9:**
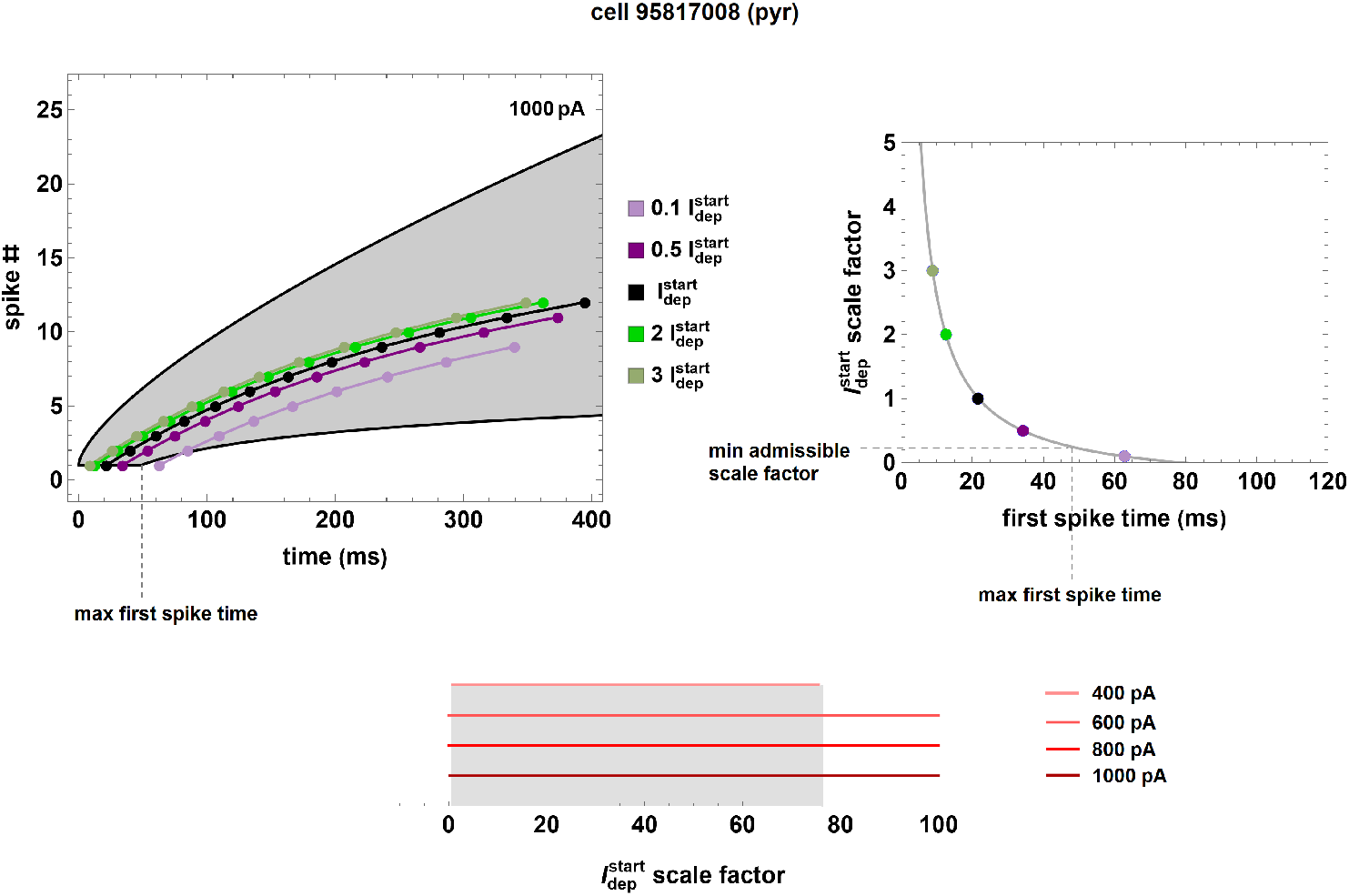
A priori validation procedure for the admissible range of the scale factors for 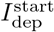. Top: (left) spike number as a function of the spike times at a current of 1000 pA for the pyramidal cell 95817008 (black curves and dots) and for 4 neuron copies (colored curves and dots) obtained by fixing the numerical values of all parameters except 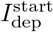; (right) determination of the range of admissible scale factors for 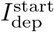 for the stimulation current of 100pA. Bottom: determination of the range of admissible scale factors for 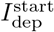 valid for all stimulation currents.

## Acknowledgments

We thank Vittorio De Falco for the support in the initial stages of the code implementation.

## Code Availability

All model and simulation files will be available in the live papers section of EBRAINS (https://live-papers.brainsimulation.eu/).

## Author contributions

AM and MM contributed to conception and design of the study. AM and AI developed the theoretical formalism and performed the mathematical analysis. CT developed the software and performed the data analysis. All authors wrote the manuscript.

## Footnotes

1 We remark that the 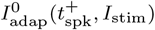 distributions for all cells provided by the optimization procedure satisfy the constraint (15). However, this constraint could be violated in determining the interpolating function (16) of each cell.

2 We recall that the perturbations of the parameter *α* automatically lead to modifying the value of the stimulation current *I*_th_ (see Section 3.1.3).

3 The parameter values of *E_L_*, *V_r_*, *V*_th_ as well as the spike times at any constant currents.

